# Physics-informed neural network for predicting myocardial mechanics from in vivo physiological measurements

**DOI:** 10.64898/2026.10.16.756787

**Authors:** Rana Raza Mehdi, Nikhil Kadivar, Sunder Neelakantan, Kyle J. Myers, Eric Wang, Melih Agraz, Peng Zhang, George Karniadakis, Gaurav Choudhary, Reza Avazmohammadi

## Abstract

Structural heart diseases (SHDs) are characterized by progressive myocardial remodeling, yet the tissue-level mechanical properties underlying this process cannot be measured directly in vivo. This limitation restricts the ability of conventional physiological measurements to fully characterize myocardial remodeling and disease progression. Here, we present a physics-informed neural network (PINN) framework that integrates constitutive biomechanics with organ-level physiological measurements to simultaneously infer myocardial biomechanical descriptors and predict physically consistent stress responses. By embedding a hyperelastic constitutive model within the learning process, the framework links physiological measurements with tissue-level mechanics while preserving biomechanical consistency. Using right ventricular remodeling in pulmonary hypertension (PH) as a representative model of SHDs, the proposed framework accurately reproduced nonlinear stress-stretch behavior and outperformed conventional data-driven machine learning approaches for stress prediction. Furthermore, incorporation of PINN-derived tissue-level descriptors with organ-level physiological measurements improved disease stratification and enhanced separation between control, mild, and severe PH remodeling states. These findings support physics-informed constitutive learning as an interpretable framework for integrating organ-level physiology with tissue-level biomechanics and establish a foundation for future biomechanics-informed characterization of myocardial remodeling across SHDs.

## 1 Introduction

Structural heart diseases (SHDs) comprise a broad and heterogeneous group of cardiac pathologies, including pressure- and volume-overload conditions, ischemic injury, congenital abnormalities, and cardiomyopathies^1^. Despite their diverse etiologies, SHDs share a common characteristic, which is progressive myocardial remodeling driven by altered mechanical loading^2^. Across disease types, this remodeling manifests as tightly coupled changes in myocardial microstructure and material behavior, including myocyte hypertrophy, collagen deposition, fiber reorientation, and increased extracellular matrix stiffness, leading to altered myocardial stiffness, anisotropy, and stress distribution^3–6^. These changes reshape the nonlinear stress–strain response of the myocardium and redistribute mechanical loads across the ventricular wall, ultimately impairing ventricular function and strongly influencing patient outcomes^7^. While global clinical metrics such as chamber volumes, ejection fraction, and pressure measurements provide indirect markers of dysfunction, they do not directly capture the tissue-level mechanical state that governs remodeling. In particular, myocardial stresses, which are fundamental descriptors of mechanical load, cannot be measured in vivo and must be inferred indirectly, highlighting a critical gap in the non-invasive characterization of tissue-level myocardial mechanics across SHDs.

Pulmonary hypertension (PH) represents a clinically important example of SHD, in which elevated pulmonary vascular resistance imposes chronic pressure overload on the right ventricle (RV)^8–11^. Under sustained overload, the RV undergoes progressive remodeling, initially through adaptive hypertrophy and subsequently through dilation, fibrosis, and eventual failure^12–14^. Importantly, RV dysfunction has emerged as a dominant determinant of outcomes in PH, often independent of pulmonary vascular resistance^15–17^. Clinical studies have demonstrated that imaging-derived indices such as RV ejection fraction, strain, and mass-to-volume ratio predict survival but still do not directly quantify tissue mechanics^18–20^. Although PH serves as a clear illustration of pressure-overload remodeling, similar biomechanical mechanisms underlie myocardial adaptation across many SHDs, including left-sided heart failure, valvular disease, and congenital conditions. Thus, PH provides a tractable model system for studying mechanically driven remodeling while maintaining relevance to broader SHD contexts.

Despite the recognized importance of myocardial mechanics in SHDs, direct characterization of tissue material properties has traditionally relied on ex-vivo mechanical testing, most commonly biaxial tensile testing of excised myocardial samples^21^. While these experiments provide ground-truth stress–strain relationships, they are inherently invasive, destructive, and limited to animal models or rare human biopsy or autopsy samples^22^. Such approaches are incompatible with longitudinal assessment, population-scale studies, or routine clinical use. Furthermore, estimating material parameters associated with myofiber and collagen mechanics generally relies on inverse modeling of ex-vivo stress-stretch measurements, which requires destructive mechanical testing for each tissue sample and limits direct application to in-vivo patient assessment and generalization across disease states^23,24^. These limitations have restricted biomechanical modeling to small, highly controlled datasets and have prevented its translation into clinically scalable tools for SHD assessment.

Data-driven methods have recently been explored to estimate myocardial mechanics from imaging and physiological measurements, but many lack explicit enforcement of physical laws, resulting in predictions that may be difficult to interpret or physically inconsistent^25–28^. An ideal framework for SHDs must therefore balance clinical accessibility, physical fidelity, and interpretability, enabling inference of tissue-level mechanical behavior from indirect in-vivo measurements while respecting the governing principles of continuum mechanics. Physics-informed neural networks (PINNs) offer a promising solution to this challenge^29^. By embedding constitutive relations, strain energy functions, and mechanical constraints directly into the learning process, PINNs enable simultaneous data fitting and physics enforcement^30,31^. In biomechanics, PINNs have been applied to estimate constitutive behavior, reconstruct stress fields, and learn hyperelastic tissue responses while enforcing the governing principles of continuum mechanics^32,33^. These properties make PINNs particularly well-suited for studying myocardial remodeling in SHDs, where tissue heterogeneity and data scarcity are common. While prior work has primarily focused on left ventricular mechanics using imaging-derived displacements^34,35^, applications to RV remodeling remain limited. In particular, physics-informed frameworks capable of predicting myocardial stress responses from clinically accessible physiological measurements while simultaneously learning subject-specific constitutive parameters for disease characterization have not been widely explored.

In this study, we propose a PINN framework for predicting myocardial stress responses and learning subject-specific constitutive parameters in SHD. The framework integrates organ-level physiological measurements with ex-vivo biaxial stress-stretch data and encodes myocardial mechanics through strain invariants derived from tissue deformation and learned in-plane material orientation. By computing stresses through automatic differentiation of a hyperelastic strain energy function, the model enforces physical consistency while simultaneously learning interpretable material parameters. PH is used as a representative SHD to evaluate whether the proposed framework can accurately predict myocardial stress responses, infer biomechanically meaningful tissue descriptors, and improve discrimination between control, mild, and severe remodeling. Although demonstrated in PH, the framework provides a foundation for extending physics-informed biomechanical characterization to other SHDs.

## 2 Materials and methods

### 2.1 Animal model and experimental design

#### 2.1.1 Animal models

All animal protocols were reviewed and approved by the Institutional Animal Care and Use Committee (IACUC) at the Providence VA Medical Center (IACUC 1633548) in accordance with the National Institutes of Health (NIH) Guide for the Care and Use of Laboratory Animals. Adult male and female Sprague-Dawley (SD) and Charles Dawley Fischer (CDF) rats were used in this study to evaluate RV remodeling in healthy and diseased states. Both sexes and multiple strains were represented in the experimental cohort. A total of 39 rats were included across three groups, including control (healthy), decompensated hypoxia (mild PH), and Sugen-hypoxia–induced PH (severe PH). The final cohort included 27 male and 12 female animals. The control group included 13 males and 6 females, the mild-PH group included 3 males and 3 females, and the severe-PH group included 11 males and 3 females. The detailed experimental procedures, including animal housing, monitoring, and euthanasia protocols, are described in our previous publication, which forms the basis of this study^5^. The current study extends this previously published work by incorporating additional samples to support the development and evaluation of the proposed PINN framework. An overview of the experimental and computational workflow is presented in Fig. 1.

**Fig. 1.**
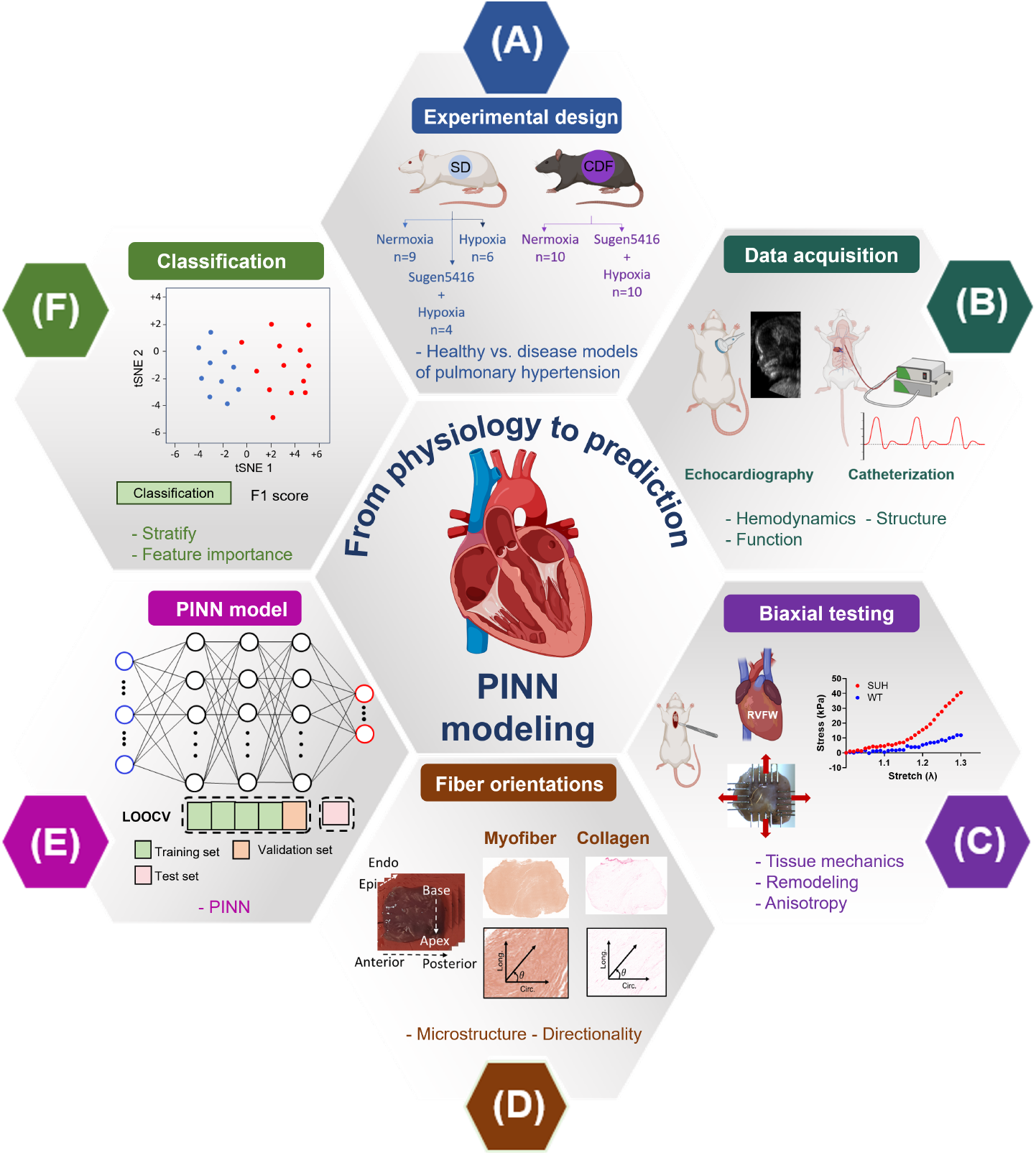
Methodology pipeline for physics-informed neural network (PINN) modeling of right ventricular (RV) remodeling. (A) Experimental design included controls (healthy), hypoxia-induced mild pulmonary hypertension (PH), and Sugen-hypoxia-induced severe PH rodent models. (B) Echocardiography and catheterization were performed to obtain hemodynamic, functional, and geometric measurements. (C) Right ventricular free-wall specimens underwent ex-vivo biaxial mechanical testing to obtain circumferential and longitudinal stress-stretch responses used for model supervision. (D) Histological analysis was performed independently to characterize myocardial microstructure; the central distribution of histologically measured myofiber orientations guided the selection of a global ±20° bound for the inferred in-plane fiber-orientation parameter, without providing specimen-specific histological inputs or supervision. (E) The PINN integrated organ-level physiological measurements and prescribed tissue stretches with a constitutive strain-energy formulation to predict myocardial stresses and infer constitutive material parameters under leave-one-out cross-validation (LOOCV). (F) Organ-level physiological features and combined organ-level plus PINN-derived tissue-level parameters were evaluated for three-class disease classification and feature-importance analysis.

#### 2.1.2 Pulmonary hypertension induction and disease grouping

PH was induced using two established protocols representing progressive stages of RV remodeling. To model mild PH, SD rats (n = 6) were placed in a hypoxia chamber (10% O_2_) for 3 weeks, inducing early vascular remodeling and compensatory RV hypertrophy. To model severe PH, a Sugen-hypoxia (SUHx) protocol was employed. CDF rats (n = 10) received a subcutaneous injection of SU5416 (20 mg/kg), a vascular endothelial growth factor receptor inhibitor, followed by 3 weeks of hypoxia (10% O_2_) and 1 week of normoxia to mimic transition into maladaptive remodeling. To increase the SUHx cohort size and capture a broader spectrum of RV remodeling, an additional four SD rats were subjected to the same SU5416 and hypoxia protocol, increasing the total SUHx sample size to 14. Normoxic controls were included to establish healthy baselines. These included SD control rats (n = 9) and CDF control rats (n = 10), which were age- and sex-matched and housed under normoxic conditions for the same duration as their respective experimental cohorts. All animal groups included both male and female rats. Animals were assigned to the control, mild PH, and severe PH groups according to the induction protocol, with disease status confirmed by hemodynamic and biomechanical assessments.

### 2.2 Data acquisition and mechanical testing

#### 2.2.1 Hemodynamic measurements (invasive and non-invasive)

Hemodynamic and echocardiographic measurements were acquired at terminal time points to comprehensively characterize RV function, as summarized in the experimental workflow (Fig. 1). Invasive catheterization was performed to obtain heart rate, right ventricular systolic pressure (RVSP), and right ventricular end-diastolic pressure (RVEDP), which served as indicators of ventricular contractile performance and relaxation. Echocardiography was used to quantify RV geometry and function, including right ventricular free-wall (RVFW) thickness, pulmonary acceleration time (PAT), tricuspid annular plane systolic excursion (TAPSE), RV systolic myocardial velocity (RV S^*′*^), and additional cardiac functional and geometric measurements. These RV-focused hemodynamic, functional, and geometric measurements were used to characterize disease severity and constituted the organ-level input feature set for the proposed PINN framework. The final model used 15 organ-level variables, including RV S^*′*^, PAT, RVFW thickness, TAPSE, heart rate, RVEDP, RVSP, systolic and diastolic diameters, systolic and diastolic volumes, stroke volume, ejection fraction, fractional shortening, and cardiac output. Additional echocardiographic measurements, including diastolic function and left ventricular outflow tract (LVOT) hemodynamics, were acquired for broader physiological characterization but were not incorporated into the PINN.

#### 2.2.2 Ex-vivo biaxial testing and histological analysis

Following terminal procedures, the RVFW tissue was harvested and immediately immersed in cold phosphate-buffered saline to preserve biomechanical integrity. The RVFW was isolated and trimmed into rectangular specimens aligned with the circumferential and longitudinal directions. The in-plane specimen dimensions varied among samples and were measured individually. Specimen thickness was measured using calipers prior to mechanical testing, and the measured specimen width perpendicular to each loading direction, together with the thickness, was used to determine the corresponding initial cross-sectional area. These specimen-specific areas were used to convert the recorded forces into first Piola-Kirchhoff stresses. Specimens were mounted in a CellScale BioTester using BioRakes and submerged in phosphate-buffered saline at approximately 22°C during testing. The testing protocol consisted of 10 preconditioning cycles followed by 10 equibiaxial loading cycles, with the unloaded specimen configuration at the beginning of the recorded loading cycle defined as the reference configuration. Stretch was calculated from the overall specimen deformation without optical-marker-based local displacement tracking. A single equibiaxial loading ratio (1:1) was applied consistently across all specimens to provide a standardized deformation protocol for cross-specimen and cross-group comparison. Measured forces and prescribed specimen deformations were converted into first Piola-Kirchhoff stress-stretch responses based on the initial specimen geometry. Stress-stretch curves were obtained for each specimen over a stretch range of up to 30%, and these measurements served as the target data for PINN training. The complete dataset comprised 39 specimens spanning control, mild PH, and severe PH groups. Histological analysis was performed independently to characterize myocardial microstructural remodeling, including myofiber and collagen orientation. Specimen-specific histological measurements were not used as PINN inputs or supervision targets; the central distribution of histologically measured myofiber orientations across the cohort was used to define a global physiologically plausible range of *±*20° for the inferred orientation parameter.

### 2.3 Constitutive modeling

For each myocardial tissue specimen, biaxial force-displacement measurements were converted into first Piola-Kirchhoff stresses in the experimentally measured circumferential and longitudinal loading directions. The corresponding stretches were denoted by *λ*_*c*_ and *λ*_*l*_, respectively. These stretches ranged from 1.0 to 1.3 during biaxial testing. Assuming incompressibility of the myocardium, the out-of-plane stretch was computed as *λ*_*n*_ = (*λ*_*c*_*λ*_*l*_)^*−*1^, thereby satisfying the incompressibility constraint *λ*_*c*_*λ*_*l*_*λ*_*n*_ = 1.

The deformation state was characterized using invariants of the right Cauchy-Green deformation tensor, **C** = **F**^*T*^ **F**, where **F** denotes the deformation gradient. The isotropic contribution was represented by the first invariant,

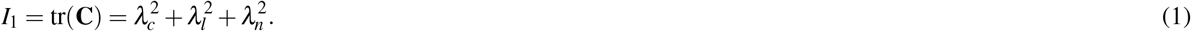

The anisotropic response was represented by the fiber, cross-fiber, and fiber-cross-fiber interaction invariants,

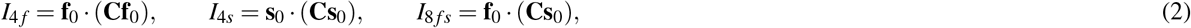

where **f**_0_ and **s**_0_ denote unit vectors defining the predominant in-plane fiber and cross-fiber material directions, respectively. The invariants *I*_4 *f*_ and *I*_4*s*_ describe deformation along these two in-plane material directions, whereas *I*_8 *fs*_ describes their interaction. Here, **s**_0_ represents an effective in-plane cross-fiber direction within the circumferential-longitudinal plane, distinct from the anatomical myocardial sheet direction associated with the three-dimensional laminar architecture. The subscript *s* is retained for consistency with the Holzapfel-Ogden notation^21^.

Although a complete three-dimensional orthotropic description of myocardium may include additional structural invariants associated with the sheet-normal direction, such as *I*_4*n*_, *I*_8 *fn*_, and *I*_8*sn*_, the present study was based on planar biaxial testing in the circumferential-longitudinal plane. Accordingly, the constitutive formulation was restricted to the in-plane invariants supported by the available deformation measurements while retaining isotropic, fiber, cross-fiber, and fiber-cross-fiber interaction contributions.

The passive myocardial response was described using the Holzapfel-Ogden hyperelastic strain-energy density function^21^,

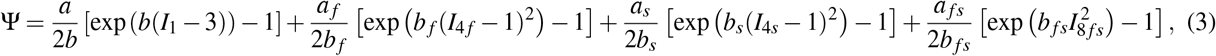

where [*a, b, a*_*f*_, *b*_*f*_, *a*_*s*_, *b*_*s*_, *a*_*fs*_, *b*_*fs*_] denote the complete set of eight constitutive parameters associated with the isotropic, fiber, cross-fiber, and fiber-cross-fiber interaction terms.

### 2.4 Physics-informed neural network (PINN) architecture

To predict myocardial stress responses while simultaneously learning constitutive material parameters, a PINN framework was developed (Fig. 2). Organ-level hemodynamic, functional, and geometric measurements were provided as inputs to a fully connected neural network comprising three hidden layers with 64, 64, and 32 neurons, respectively, and hyperbolic tangent (tanh) activation functions. For each specimen, the network inferred four constitutive magnitude parameters, (*a, a*_*f*_, *a*_*s*_, *a*_*fs*_), together with one in-plane fiber-orientation parameter, *θ*_*f*_. The material parameters were constrained to remain positive using a softplus activation function, whereas the magnitude of *θ*_*f*_ was bounded to *±* 20° about the circumferential axis, guided by the central distribution of histologically measured myofiber orientations across the cohort. This global bound was applied identically to all specimens; specimen-specific histological orientations were not provided to the network. Orthogonality of the in-plane material directions was enforced by defining *θ*_*s*_ = *θ*_*f*_ + 90°. The prescribed biaxial stretches, *λ*_*c*_ and *λ*_*l*_, were combined with the inferred in-plane fiber-orientation parameter to construct the strain invariants. Thus, *λ*_*c*_ and *λ*_*l*_ describe deformation along the experimental loading axes, whereas *θ*_*f*_ defines the orientation of the material fiber direction relative to those axes. Although the constitutive formulation in Eq. 3 contains eight material parameters, simultaneous estimation of the constitutive magnitude parameters and their associated exponential coefficients can introduce parameter-identifiability challenges, particularly with a limited set of biaxial loading conditions. Therefore, the nonlinear exponential coefficients were fixed as *b* = *b*_*f*_ = *b*_*s*_ = *b*_*fs*_ = 1, consistent with our previous characterization of RV myocardial mechanics in PH^5^, while the four constitutive magnitude parameters (*a, a*_*f*_, *a*_*s*_, *a*_*fs*_) and the in-plane fiber-orientation parameter *θ*_*f*_ were inferred by the network. The same fixed coefficients were applied to all specimens to provide a common constitutive representation across strains and disease groups while reducing the number of simultaneously identifiable parameters. The inferred *θ*_*f*_ parameterized the in-plane fiber direction, **f**_0_ = [cos(*θ*_*f*_), sin(*θ*_*f*_), 0]^*T*^, while the orthogonal cross-fiber direction **s**_0_ was defined using *θ*_*s*_ = *θ*_*f*_ + 90°. These directions, together with the prescribed circumferential and longitudinal stretches, were used to evaluate the constitutive invariants, which were subsequently incorporated into the strain-energy density function (Eq. 3). Within the differentiable constitutive framework, the first Piola-Kirchhoff stresses in the circumferential and longitudinal directions were obtained through automatic differentiation of the strain-energy density function with respect to the prescribed stretches,

**Fig. 2.**
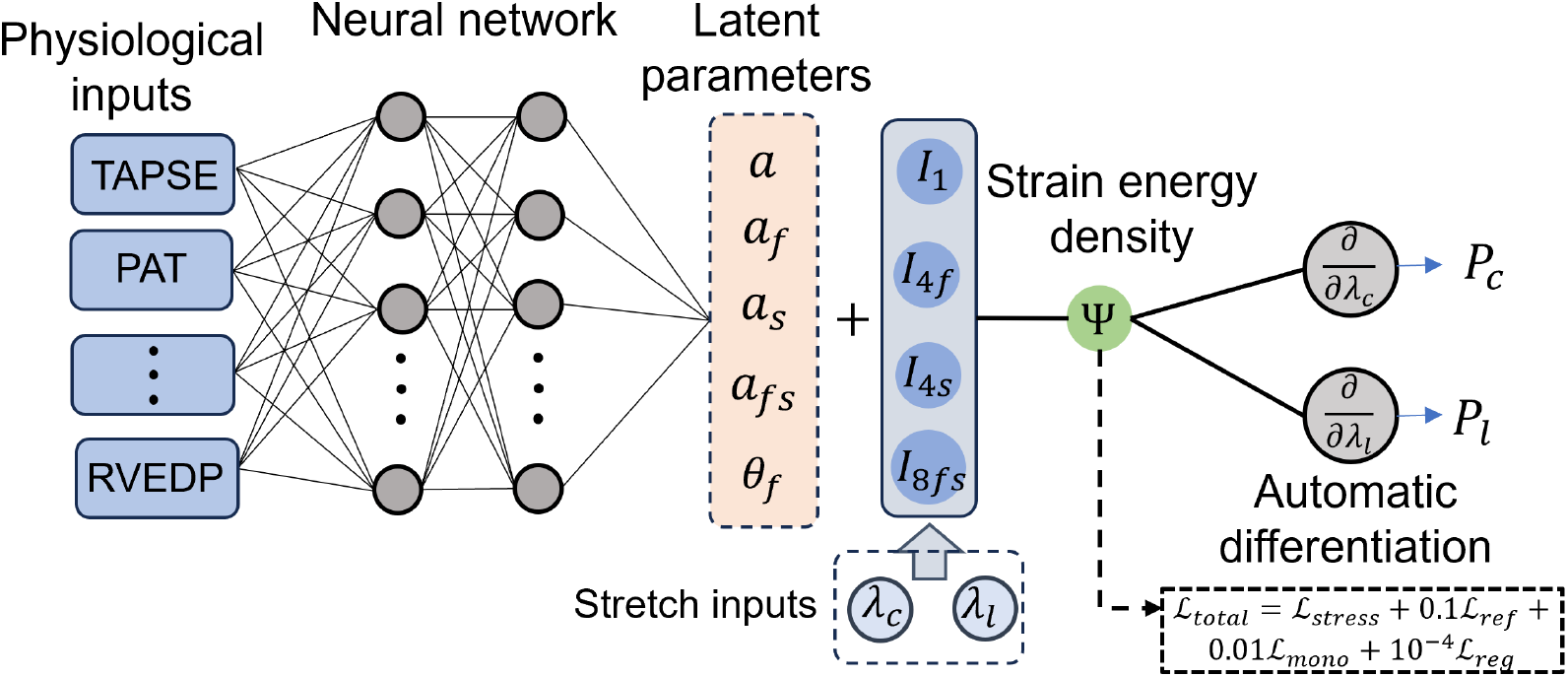
Architecture of the proposed physics-informed neural network (PINN) for prediction of myocardial stress responses and inference of constitutive biomechanical parameters. Hemodynamic, functional, and geometric measurements were provided as inputs to a fully connected neural network, which inferred four constitutive magnitude parameters and one in-plane fiber-orientation parameter, *θ*_*f*_, for each specimen. The orthogonal cross-fiber direction was defined by *θ*_*s*_ = *θ*_*f*_ + 90°. The inferred in-plane fiber-orientation parameter was combined with the prescribed circumferential and longitudinal stretches to construct the constitutive invariants, which were subsequently used together with the inferred constitutive magnitude parameters to evaluate the hyperelastic strain-energy density function, Ψ. Circumferential and longitudinal first Piola-Kirchhoff stresses were obtained through automatic differentiation of Ψ. Model training was supervised using experimentally measured stress-stretch data, with reference-state, monotonicity, and parameter-magnitude terms included as numerical regularizers.

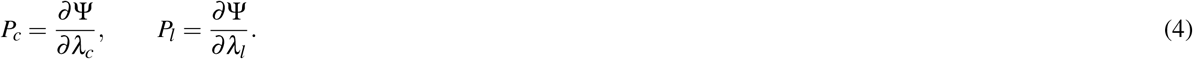

The primary data-fidelity term minimized the mean squared error between the predicted and experimentally measured stresses in the circumferential and longitudinal directions,

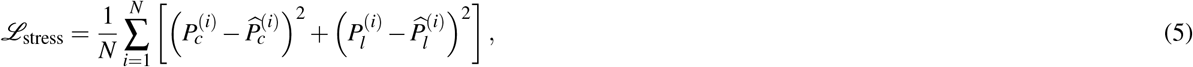

where *N* denotes the total number of stress-stretch observations, and 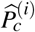 and 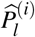 represent the experimentally measured first Piola-Kirchhoff stresses.

As an auxiliary numerical regularizer, a reference-state loss penalized residual strain energy and stresses at *λ*_*c*_ = *λ*_*l*_ = 1,

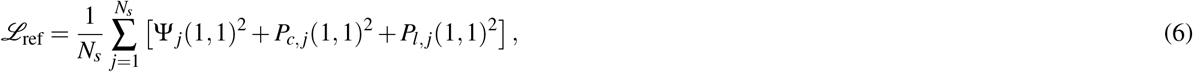

where *N*_*s*_ denotes the number of specimens in the training batch.

A second auxiliary regularization term penalized negative tangent stiffness values during optimization,

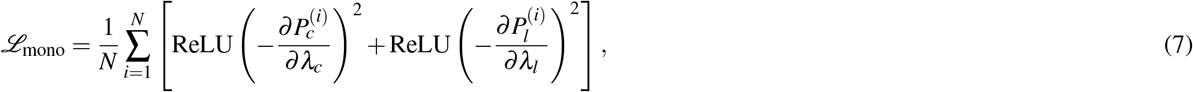

where ReLU(*x*) = max(0, *x*) penalizes only negative stress gradients and is used solely as a one-sided penalty function. Finally, a parameter regularization term was introduced to discourage unrealistically large constitutive parameters,

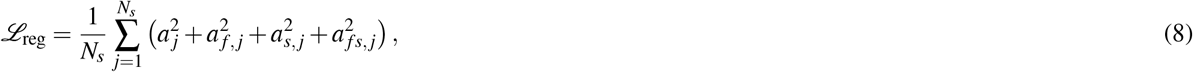

and the overall training objective was defined as

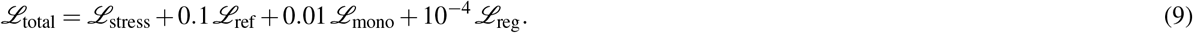

This composite objective prioritized agreement with the experimentally measured biaxial stress-stretch responses while retaining the reference-state, monotonicity, and parameter-magnitude terms as auxiliary numerical regularizers rather than independent physical constraints.

### 2.5 Model training and evaluation

A leave-one-out cross-validation (LOOCV) strategy was employed to maximize the use of the limited experimental dataset while providing an out-of-sample assessment of model generalization. For each fold, one of the 39 myocardial specimens was held out for testing, while the remaining 38 specimens were used for model development. From the training set, 20% of the specimens were randomly allocated to a validation set for model selection, and the remaining specimens were used for optimization. All data partitioning was performed at the specimen level; multiple stress-stretch observations from the same specimen were used as within-specimen mechanical supervision and were not split across training, validation, and test sets. The PINN was trained using the Adam optimizer with early stopping based on the validation loss. The model corresponding to the lowest validation loss was retained for testing. This procedure was repeated until every specimen had served once as the independent test sample. For each test specimen, the trained PINN predicted the constitutive material parameters, in-plane fiber-orientation parameter, and corresponding biaxial stress responses across the prescribed stretch range. Model performance was evaluated using the coefficient of determination (*R*^2^) and root mean squared error (RMSE) between the predicted and experimentally measured first Piola-Kirchhoff stresses. Predictions from all LOOCV folds were aggregated to quantify overall predictive performance and were subsequently used for statistical analyses and disease classification. An additional sensitivity analysis comparing Adam and self-scaled Broyden (SSBroyden) optimization under a fixed-orientation configuration is described in Supplementary Data S1.4.

### 2.6 Comparison with conventional machine learning models

To benchmark the predictive performance of the proposed PINN, three conventional machine learning (ML) regression models, including a multilayer perceptron (MLP), random forest (RF), and extreme gradient boosting (XGBoost), were trained to predict biaxial stress responses using the same organ-level physiological inputs and experimentally prescribed stretch measurements as the PINN. All baseline models were trained and evaluated using the identical LOOCV protocol described above to ensure a fair comparison. Model performance was quantified using the *R*^2^ and RMSE between the predicted and experimentally measured first Piola-Kirchhoff stresses for each specimen.

### 2.7 Conventional inverse optimization of constitutive parameters

To provide a conventional reference for the inferred in-plane fiber-orientation parameter, specimen-specific nonlinear least-squares optimization was performed using the same reduced Holzapfel–Ogden constitutive formulation. For each specimen, the four constitutive magnitude parameters (*a, a*_*f*_, *a*_*s*_, *a*_*fs*_) and the in-plane fiber-orientation parameter *θ*_*f*_ were estimated by directly fitting the experimentally measured circumferential and longitudinal stress–stretch responses. The nonlinear exponential coefficients were fixed at *b* = *b*_*f*_ = *b*_*s*_ = *b*_*fs*_ = 1, consistent with the PINN formulation. The orientation parameter was constrained within the same global *±*20° range used in the PINN, with the cross-fiber direction defined orthogonally. The inverse-optimized orientation estimates were compared with the PINN-inferred orientation estimates (Supplementary Fig. S4).

### 2.8 Disease classification using organ- and tissue-level features

Beyond stress prediction, the potential of PINN-derived tissue-level biomechanical parameters to improve discrimination of disease severity beyond conventional organ-level measurements was evaluated. Two complementary feature representations were constructed for this purpose. The first consisted of the organ-level hemodynamic, functional, and geometric measurements used as PINN inputs, whereas the second augmented these measurements with the specimen-specific constitutive and orientation parameters inferred by the PINN. A multinomial logistic regression classifier was trained to distinguish *control, mild PH*, and *severe PH* specimens using a LOOCV framework. Classification performance was evaluated using precision, recall, F1-score, overall accuracy, and three-class confusion matrices. To qualitatively assess class separability, t-distributed stochastic neighbor embedding (t-SNE) was independently applied to the organ-level and combined organ + tissue-level feature spaces for two-dimensional visualization. To further characterize group separation in the two-dimensional t-SNE representations, pairwise Euclidean distances between disease-group centroids were calculated in the respective reduced feature spaces, with larger distances indicating greater separation within each visualization. Finally, feature importance was assessed from the trained multinomial logistic regression models to identify the organ-level and tissue-level variables contributing most strongly to disease classification. Prior to model training, all features were standardized to zero mean and unit variance. Feature importance was quantified using the absolute values of the learned regression coefficients, averaged across classes and LOOCV folds, with larger values indicating greater contributions to disease discrimination.

### 2.9 Statistical analysis

Statistical analyses were performed using GraphPad Prism. Group-wise comparisons of PINN-inferred constitutive parameters were performed separately for each parameter using ordinary one-way analysis of variance (ANOVA) followed by Dunnett’s multiple-comparisons test, with the control group used as the reference. Physiological and geometric measurements reported in Table 1 were analyzed using ordinary two-way ANOVA followed by Tukey’s multiple-comparisons test for Control versus PH comparisons within each strain. Additional physiological measurements reported in Supplementary Table S1 were compared between pooled Control and PH groups using a two-tailed unpaired Welch’s *t*-test. Statistical significance was defined as *p <* 0.05.

**Table 1.** Hemodynamic, echocardiographic, and geometric measurements for the control and pulmonary hypertension (PH) groups in Sprague-Dawley and Fischer rat strains. Hemodynamic measurements include heart rate, right ventricular end-diastolic pressure (RVEDP), and right ventricular systolic pressure (RVSP). Echocardiographic measurements include right ventricular systolic myocardial velocity (RV S^*′*^), pulmonary acceleration time (PAT), tricuspid annular plane systolic excursion (TAPSE), cardiac functional indices, and right ventricular geometric measurements. The PH group comprises both mild (hypoxia-induced) and severe (Sugen-hypoxia-induced) PH models. Values are reported as mean *±*s tandard deviation. Adjusted *p*-values are from Tukey’s multiple comparisons test following two-way ANOVA. Additional echocardiographic measurements are provided in Supplementary Table S1, and histological orientation measurements are presented in Supplementary Fig. S4.

| Hemodynamic measurements |  |  |  |  |
| --- | --- | --- | --- | --- |
| | Strain | Control | PH | $p$-value |
| Heart Rate (BPM) | Sprague-Dawley | 378.44 ± 41.72 | 363.14 ± 26.38 | 0.8102 |
| | Fischer | 371.67 ± 30.22 | 348.72 ± 16.69 | 0.4481 |
| RVEDP (mmHg) | Sprague-Dawley | 6.15 ± 0.92 | 8.03 ± 1.67 | 0.2818 |
| | Fischer | 6.36 ± 0.73 | 9.56 ± 3.05 | 0.0038 |
| RVSP (mmHg) | Sprague-Dawley | 28.47 ± 5.60 | 56.86 ± 15.01 | 0.0022 |
| | Fischer | 27.28 ± 3.69 | 97.02 ± 21.89 | < 0.0001 |
| Echocardiographic functional indices |  |  |  |  |
| RV S' (mm/s) | Sprague-Dawley | 43.73 ± 9.65 | 33.67 ± 10.87 | 0.1396 |
| | Fischer | 38.78 ± 8.28 | 30.75 ± 4.89 | 0.2889 |
| PAT (ms) | Sprague-Dawley | 33.87 ± 2.67 | 24.39 ± 7.18 | 0.0030 |
| | Fischer | 25.87 ± 4.81 | 13.68 ± 1.44 | < 0.0001 |
| TAPSE (mm) | Sprague-Dawley | 1.78 ± 0.78 | 1.12 ± 0.51 | 0.4040 |
| | Fischer | 1.89 ± 0.79 | 0.97 ± 0.26 | 0.0192 |
| Fractional Shortening (%) | Sprague-Dawley | 54.33 ± 8.73 | 54.53 ± 8.36 | 0.9999 |
| | Fischer | 60.55 ± 8.78 | 70.62 ± 22.96 | 0.4325 |
| Ejection Fraction (%) | Sprague-Dawley | 83.06 ± 6.35 | 83.85 ± 7.01 | 0.9883 |
| | Fischer | 88.29 ± 5.81 | 89.69 ± 11.20 | 0.9824 |
| Cardiac Output (mL/min) | Sprague-Dawley | 72.91 ± 15.99 | 46.64 ± 13.37 | 0.0157 |
| | Fischer | 53.12 ± 23.40 | 20.74 ± 9.26 | 0.0013 |
| Geometric measurements |  |  |  |  |
| RVFW thickness (echocardiography) (mm) | Sprague-Dawley | 0.54 ± 0.13 | 1.37 ± 0.68 | 0.0005 |
| | Fischer | 0.54 ± 0.16 | 1.32 ± 0.24 | 0.0010 |
| End-systolic diameter (mm) | Sprague-Dawley | 3.12 ± 0.79 | 2.57 ± 0.72 | 0.5745 |
| | Fischer | 2.25 ± 0.79 | 1.29 ± 1.14 | 0.1339 |
| End-diastolic diameter (mm) | Sprague-Dawley | 6.72 ± 0.88 | 5.55 ± 0.72 | 0.0821 |
| | Fischer | 5.59 ± 1.19 | 3.85 ± 0.97 | 0.0035 |
| End-systolic volume (μL) | Sprague-Dawley | 42.52 ± 19.05 | 27.21 ± 16.73 | 0.2245 |
| | Fischer | 21.09 ± 14.29 | 10.71 ± 15.26 | 0.5701 |
| End-diastolic volume (μL) | Sprague-Dawley | 238.78 ± 64.48 | 154.59 ± 46.02 | 0.0224 |
| | Fischer | 163.28 ± 70.91 | 70.31 ± 38.45 | 0.0093 |
| Stroke volume (μL) | Sprague-Dawley | 196.26 ± 49.76 | 127.39 ± 31.65 | 0.0162 |
| | Fischer | 142.19 ± 59.79 | 59.59 ± 26.82 | 0.0026 |

## 3 Results

### 3.1 Physiological characterization of pulmonary hypertension

Hemodynamic, echocardiographic, and geometric measurements demonstrated PH-associated physiological alterations (Table 1). Relative to their strain-matched controls, PH animals exhibited elevated right ventricular systolic pressure (RVSP), with significant differences observed in both Sprague-Dawley and Fischer cohorts. Right ventricular end-diastolic pressure (RVEDP) was also elevated, with a significant difference in the Fischer cohort. Pulmonary acceleration time (PAT) decreased with PH, whereas RVFW thickness increased, consistent with increased pulmonary vascular load and RV hypertrophic remodeling. Cardiac output was significantly reduced in both strains. Several additional functional and geometric indices also differed between control and PH animals, whereas global systolic measures such as ejection fraction and fractional shortening remained largely preserved. Additional echocardiographic and LVOT measurements are summarized in Supplementary Table S1. Together, these measurements characterized the hemodynamic and structural remodeling represented in the experimental cohort.

### 3.2 Biaxial mechanical characterization

Representative biaxial stress-stretch responses of RVFW specimens from the control, mild PH, and severe PH groups are shown in Fig. 3. Progressive changes in myocardial mechanical behavior were observed with increasing disease severity. Control specimens exhibited the most compliant response in both the circumferential and longitudinal directions, whereas RVFW tissues from mild PH animals demonstrated increased stresses over the same stretch range. The severe PH group exhibited the steepest stress-stretch responses, indicating the greatest tissue stiffness. The nonlinear mechanical response became increasingly pronounced from the control to the severe PH groups, consistent with progressive stiffening of the RV myocardium. Although inter-specimen variability increased in the diseased groups, particularly at higher stretch levels, the overall trend of increasing stiffness with disease severity was preserved in both loading directions. These experimentally measured biaxial stress-stretch responses served as the ground truth for subsequent PINN training and evaluation.

**Fig. 3.**
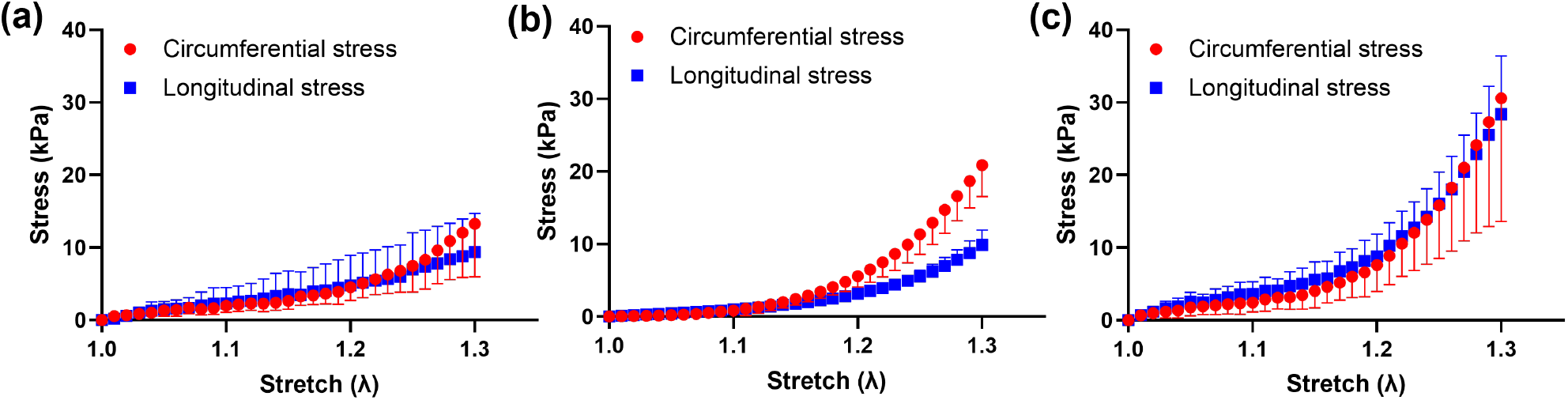
Biaxial stress–stretch responses of right ventricular free wall (RVFW) tissue across groups. Experimental circumferential and longitudinal stresses are shown for (a) control (n=19), (b) mild pulmonary hypertension (PH) (decompensated hypoxia, n=6), and (c) severe PH (Sugen-hypoxia–induced, n=14). Error bars represent the variability across specimens within each group. Progressive stiffening with increasing disease severity is evident in both loading directions.

### 3.3 PINN prediction of biaxial stress responses

Representative PINN predictions for the control, mild PH, and severe PH groups are shown in Fig. 4. Across all disease groups, the predicted circumferential and longitudinal first Piola-Kirchhoff stresses closely matched the experimentally measured biaxial stress-stretch responses, demonstrating that the proposed framework accurately captured the nonlinear mechanical behavior of the RVFW. Close agreement was observed throughout the prescribed stretch range, with the predicted curves reproducing both the magnitude and characteristic shape of the experimental responses for healthy and remodeled myocardium.

**Fig. 4.**
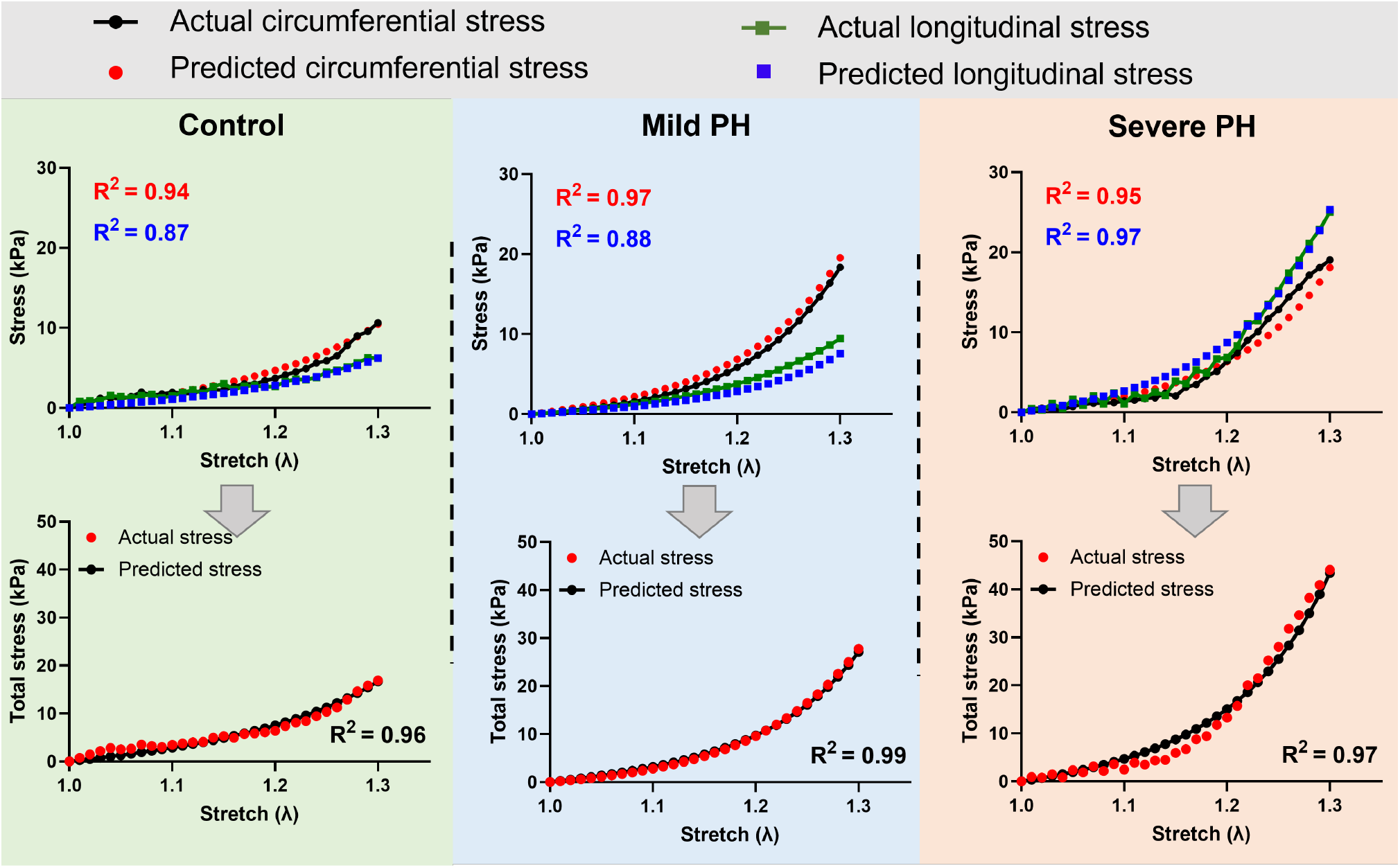
Representative stress predictions for the control, mild pulmonary hypertension (PH), and severe PH groups using the proposed physics-informed neural network (PINN). The upper row compares the experimentally measured and predicted circumferential and longitudinal first Piola-Kirchhoff stresses over the prescribed stretch range for representative specimens from each disease group. The lower row compares the corresponding experimentally measured and predicted total stress, defined as the sum of the circumferential and longitudinal stress components. The coefficient of determination (*R*^2^) reported in each panel quantifies the agreement between the predicted and experimentally measured stress responses.

The predictive performance of the proposed PINN was further compared with three conventional ML regression models, including MLP, RF, and XGBoost (Fig. 5). Across all experimental groups, the PINN achieved consistently higher *R*^2^ values than the baseline ML approaches. Although all models performed well for some specimens, the conventional ML models exhibited larger variability and produced negative *R*^2^ values for a subset of control and severe PH specimens, indicating predictions that were less accurate than a mean-response baseline. In contrast, the PINN maintained positive predictive performance across all disease groups. Consistent with the *R*^2^ results, the RMSE analysis showed lower prediction errors for the PINN than for the conventional ML models across the control, mild PH, and severe PH groups (Supplementary Fig. S3).

**Fig. 5.**
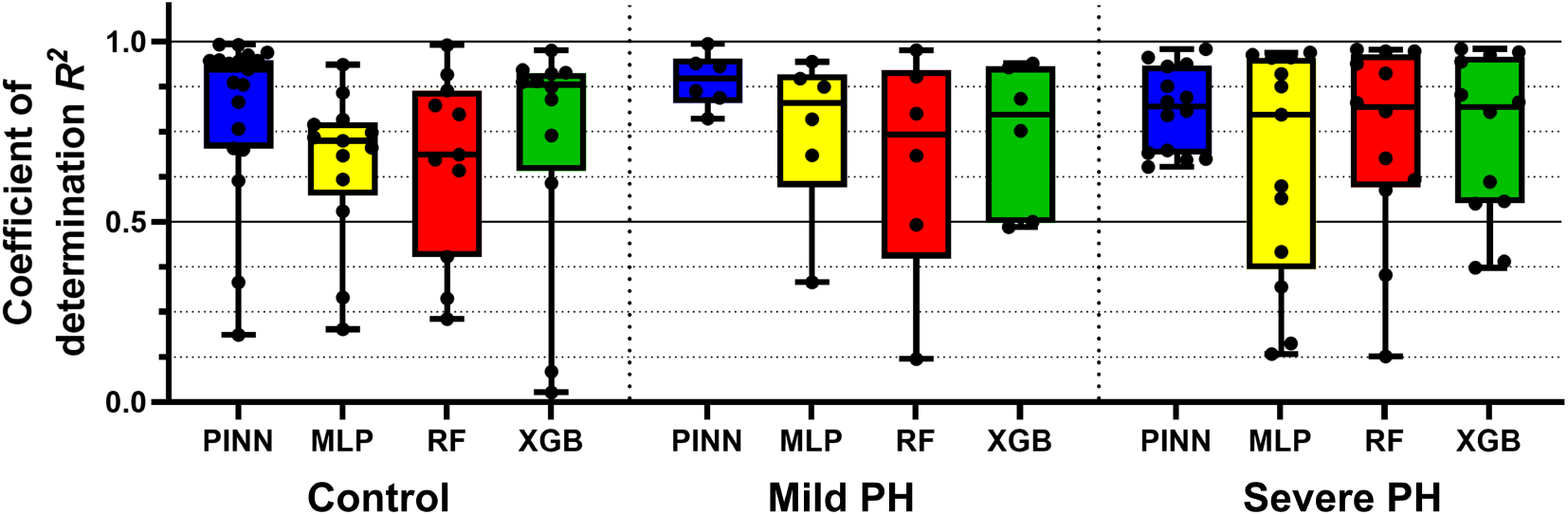
Comparison of stress-prediction performance across different modeling approaches. Negative *R*^2^ values, which indicate performance worse than a mean-response baseline, were omitted from the displayed range for visualization clarity. These omitted negative values occurred for the multilayer perceptron, random forest, and extreme gradient boosting models in the control and severe pulmonary hypertension groups.

To further evaluate prediction accuracy throughout the loading protocol, differences between the predicted and experimentally measured total stresses were evaluated at representative mid- and maximum-stretch levels (Supplementary Fig. S2). Prediction differences were smaller at intermediate stretch and increased near the maximum applied stretch, where the nonlinear mechanical response became more pronounced. Training convergence of the proposed PINN, including the evolution of the training and validation losses and corresponding *R*^2^ values, is provided in Supplementary Fig. S1. An additional optimizer comparison between Adam and SSBroyden is presented in Supplementary Fig. S5 and demonstrated broadly comparable stress agreement between the two optimization strategies.

### 3.4 PINN-inferred constitutive parameters

The specimen-specific constitutive parameters inferred by the PINN are summarized in Fig. 6. The reduced constitutive formulation enabled estimation of the isotropic (*a*), fiber (*a*_*f*_), cross-fiber (*a*_*s*_), and fiber-cross-fiber interaction (*a*_*fs*_) stiffness parameters from the organ-level physiological measurements within the physics-informed framework. Group-wise comparisons were performed between the control, mild PH, and severe PH cohorts to evaluate disease-associated changes in the inferred biomechanical parameters. The inferred parameter distributions demonstrated remodeling-associated differences across disease groups. Significant pairwise differences were observed for the isotropic parameter *a*, fiber stiffness parameter *a*_*f*_, and cross-fiber stiffness parameter *a*_*s*_, whereas the fiber-cross-fiber interaction parameter *a*_*fs*_ exhibited greater overlap among groups (Fig. 6). These findings indicate that the PINN-derived constitutive parameters captured changes in myocardial mechanical behavior associated with progressive RV remodeling. The corresponding PINN-inferred orientation parameter is presented in Supplementary Fig. S4. The inferred in-plane fiber-orientation parameter also exhibited disease-associated differences among the experimental groups, while the cross-fiber direction was defined by the imposed orthogonality relationship. Together, the constitutive and orientation parameters provided tissue-level biomechanical descriptors that were subsequently combined with organ-level physiological measurements for disease classification.

**Fig. 6.**
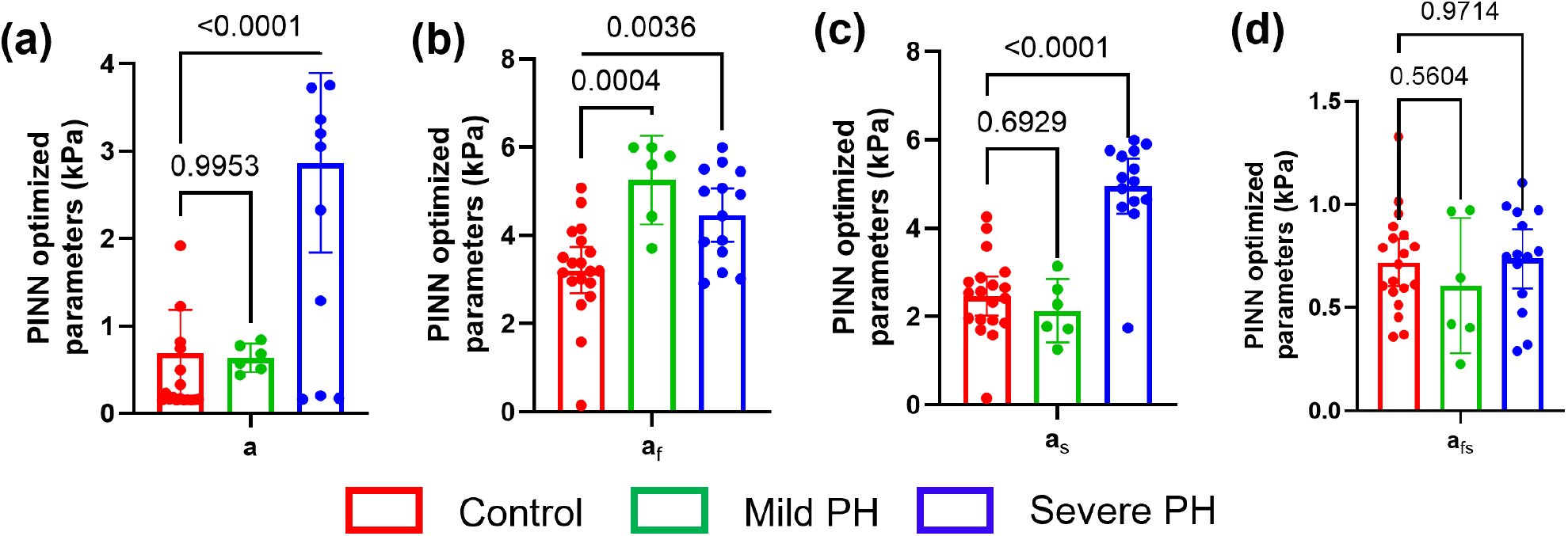
PINN-inferred constitutive material parameters across the control, mild pulmonary hypertension (PH), and severe PH groups. Group-wise distributions of the isotropic stiffness parameter (*a*), fiber stiffness parameter (*a*_*f*_), cross-fiber stiffness parameter (*a*_*s*_), and fiber-cross-fiber interaction parameter (*a*_*fs*_) are shown. Error bars represent the 95% confidence intervals across specimens within each group. Statistical comparisons were performed using ordinary one-way analysis of variance (ANOVA) followed by Dunnett’s multiple-comparisons test, with the control group as the reference; adjusted *p*-values are indicated above each panel. Corresponding PINN-inferred and conventional inverse-optimized orientation estimates are presented in Supplementary Fig. S4.

### 3.5 Disease classification using organ- and tissue-level features

Disease classification performance was evaluated using multinomial logistic regression models trained with either organ-level physiological measurements alone or a combined feature set incorporating both organ-level measurements and PINN-derived tissue-level parameters. Classification performance was assessed using LOOCV and quantified using precision, recall, F1-score, and overall accuracy (Table 2). Using organ-level measurements alone, the classifier achieved an overall accuracy of 0.87. All control and severe PH specimens were correctly classified, whereas five of the six mild PH specimens were misclassified as control (Fig. 7a). Incorporation of the PINN-derived constitutive and orientation parameters increased the overall accuracy to 0.97 and reduced the mild-PH misclassifications to one specimen (Fig. 7b). The largest improvement was observed for the mild PH group, for which recall increased from 0.17 to 0.83 and F1-score increased from 0.29 to 0.91 (Table 2). These results indicate that the PINN-derived tissue-level parameters improved discrimination of the intermediate remodeling state.

**Table 2.** Classification performance of multinomial logistic regression models trained using (i) organ-level physiological measurements alone and (ii) combined organ-level and PINN-derived tissue-level biomechanical parameters. Tissue-level features consisted of PINN-inferred constitutive and orientation parameters. Performance was evaluated using leave-one-out cross-validation and summarized by class-specific precision, recall, and F1-score together with overall model accuracy.

| Trained on organ-level physiological features |  |  |  |  |
| --- | --- | --- | --- | --- |
|  | Precision | Recall | F1-score | Accuracy |
| Control | 0.79 | 1.00 | 0.88 | 0.87 |
| Mild PH | 1.00 | 0.17 | 0.29 |  |
| Severe PH | 1.00 | 1.00 | 1.00 |  |
| Trained on organ-level + PINN-derived biomechanical features |  |  |  |  |
| Control | 0.95 | 1.00 | 0.97 | 0.97 |
| Mild PH | 1.00 | 0.83 | 0.91 |  |
| Severe PH | 1.00 | 1.00 | 1.00 |  |

**Fig. 7.**
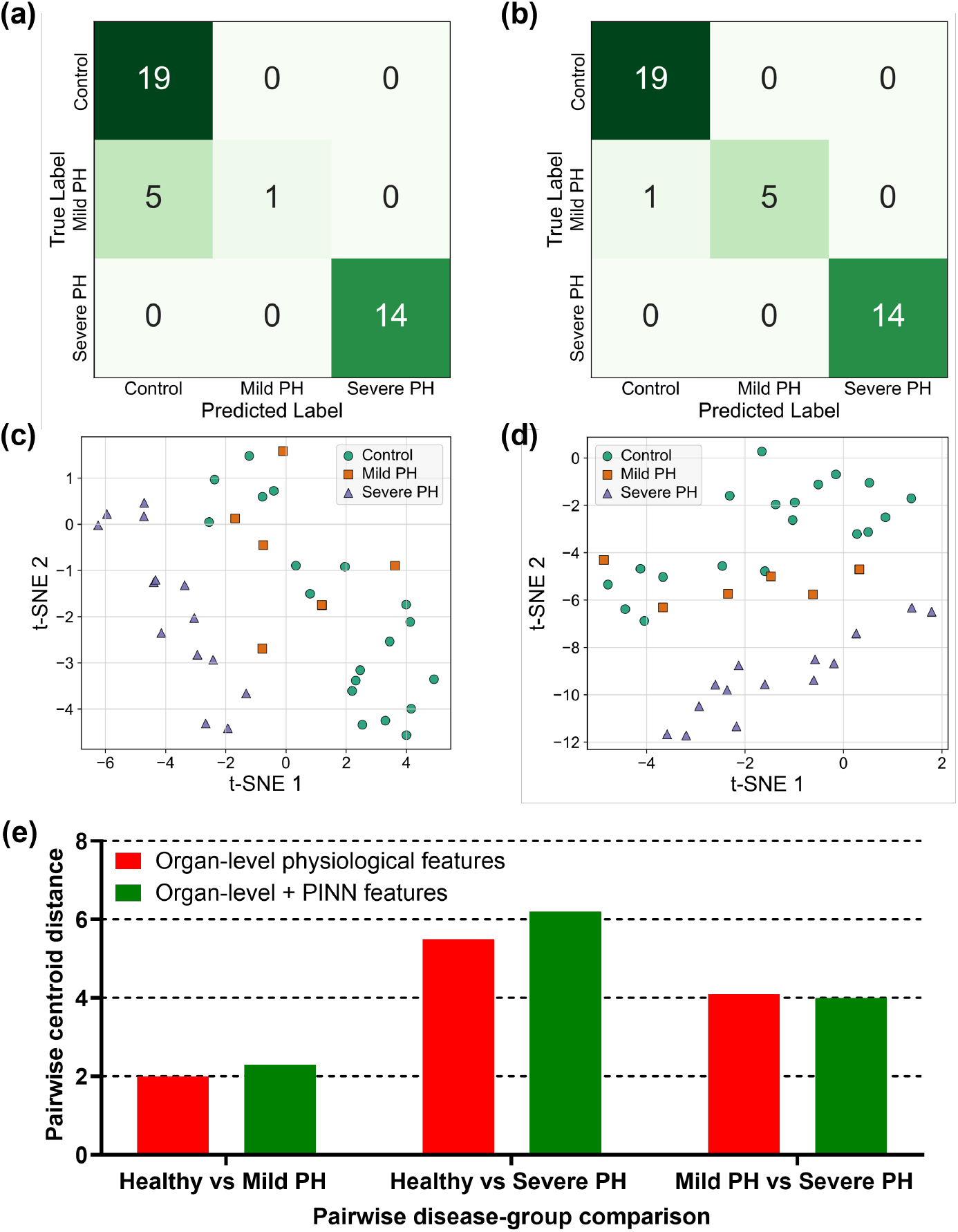
Disease classification using organ-level and PINN-derived tissue-level biomechanical features. (a,b) Confusion matrices obtained using multinomial logistic regression trained with organ-level physiological measurements alone and with combined organ-level and PINN-derived tissue-level biomechanical features, respectively. (c,d) Corresponding two-dimensional t-SNE visualizations of the organ-level and combined organ-level plus PINN-derived tissue-level feature representations, respectively. (e) Pairwise centroid distances were computed within the respective two-dimensional t-SNE embeddings for the organ-level and combined organ-level plus PINN-derived tissue-level feature representations. Larger centroid distances indicate greater separation between disease groups within each visualization.

To further examine feature-space organization, t-SNE was applied separately to the organ-level and combined organ + tissue-level feature sets (Fig. 7c-d). The severe PH group remained well separated under both feature representations, whereas overlap between the control and mild PH groups was reduced after incorporation of the PINN-derived parameters. Consistent with these qualitative observations, pairwise centroid distances calculated within the respective two-dimensional t-SNE embeddings showed modest changes between the organ-level and combined feature representations (Fig. 7e). Thus, the improvement in classification performance was more pronounced than the change in global centroid separation.

### 3.6 Feature importance analysis

Coefficient-based feature importance was used to identify the variables contributing most strongly to disease classification (Fig. 8). In the organ-level model, RVFW thickness and RVSP were the highest-ranked features, followed by PAT and RVEDP. TAPSE, heart rate, cardiac output, fractional shortening, systolic volume, and diastolic volume showed comparatively smaller contributions. Following incorporation of the PINN-derived tissue-level parameters, the inferred in-plane fiber-orientation parameter *θ*_*f*_ became the highest-ranked feature, followed by RVFW thickness and the PINN-inferred fiber stiffness parameter *a*_*f*_. RVSP and the cross-fiber stiffness parameter *a*_*s*_ also ranked among the most influential variables, followed by RVEDP, PAT, and the isotropic stiffness parameter *a*. The presence of both organ-level physiological measurements and PINN-derived biomechanical parameters among the highest-ranked features is consistent with the improved classification performance obtained using the combined feature representation. Together, these results indicate that the combined classifier incorporated information related to RV pressure loading and geometric remodeling together with PINN-derived descriptors of myocardial material behavior and in-plane orientation.

**Fig. 8.**
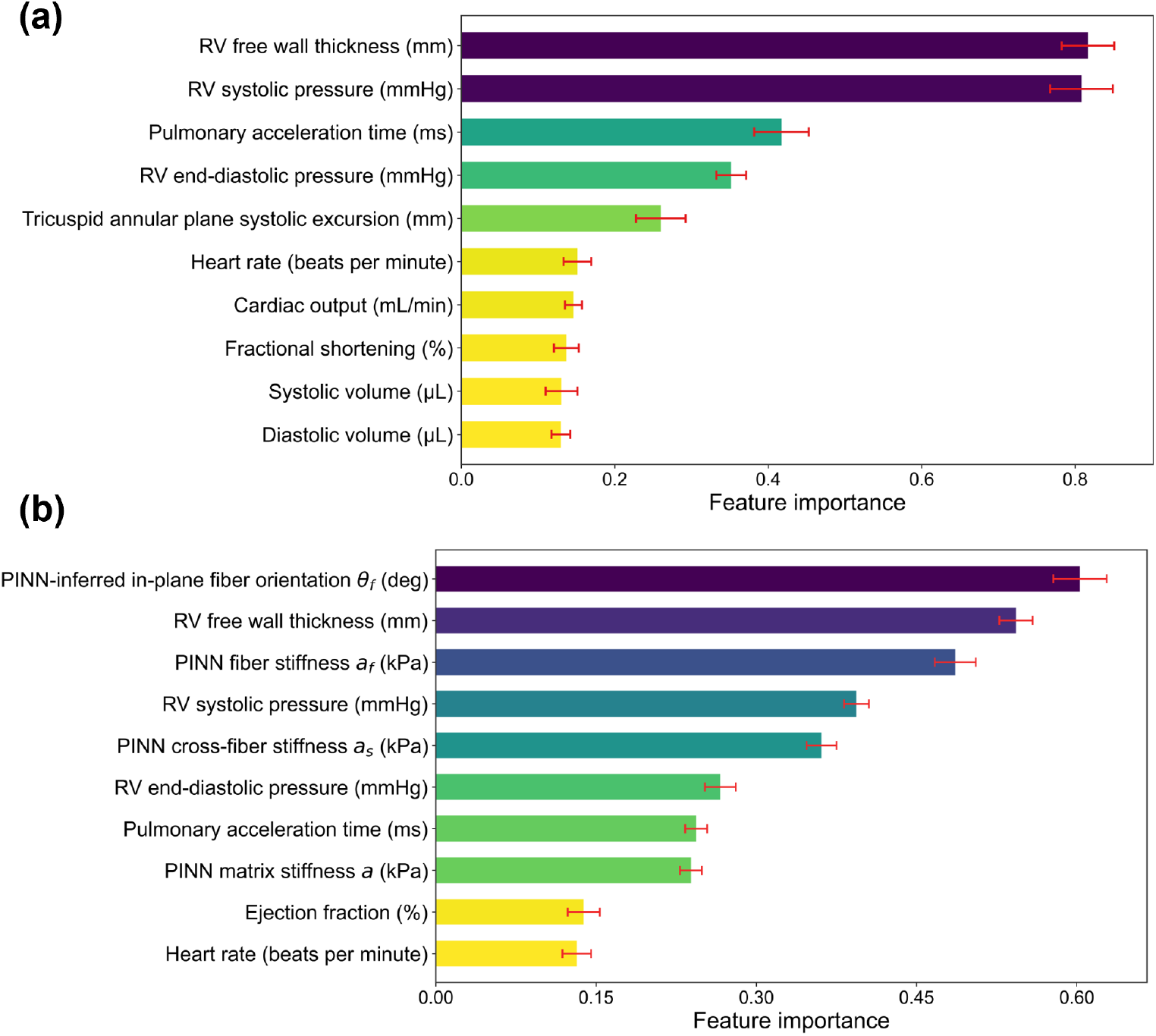
Feature importance derived from multinomial logistic regression models trained using (a) organ-level physiological measurements alone and (b) combined organ-level and PINN-derived tissue-level biomechanical parameters. Tissue-level features consisted of PINN-inferred constitutive and orientation parameters. Feature importance was quantified using the absolute values of the standardized logistic regression coefficients, averaged across classes and leave-one-out cross-validation (LOOCV) folds, with larger values indicating greater contributions to disease classification. Error bars represent *±*1 standard deviation of feature importance across LOOCV folds. The features are ordered by decreasing importance within each model.

## 4 Discussion

### 4.1 Physics-informed learning accurately predicted RV biomechanical behavior

The proposed PINN accurately reproduced the nonlinear biaxial stress-stretch behavior of RV myocardium across control, mild PH, and severe PH specimens. Representative predictions demonstrated close agreement with the experimental responses (Fig. 4), while quantitative comparisons across all specimens showed consistently high prediction accuracy and improved performance relative to conventional ML models (Fig. 5). The improved predictive performance of the PINN relative to the conventional ML baselines is likely related to the constitutive physics embedded within the learning framework. Whereas the MLP, RF, and XGBoost models learn a direct mapping from the organ-level measurements and prescribed stretches to stress, the PINN infers a reduced set of constitutive and orientation parameters whose corresponding stress-stretch response is generated through the embedded Holzapfel-Ogden formulation. This mechanically constrained representation restricts the admissible solution space and provides a physics-based inductive bias, which may be particularly beneficial for learning from the relatively small experimental cohort. Consequently, the predicted stresses remain directly linked to the underlying constitutive relationships rather than relying solely on statistical correlations between inputs and outputs. The reference-state and monotonicity terms were retained as auxiliary numerical regularizers within the physics-informed training objective.

Prediction accuracy remained high across all remodeling stages despite the greater mechanical heterogeneity observed in the diseased myocardium. Comparison of the predicted and experimental stresses at representative loading levels (Supplementary Fig. S2) showed smaller differences at the representative intermediate stretch than at the maximum stretch, indicating closer agreement at the intermediate loading level. The larger discrepancies observed near the highest stretch levels likely reflect the increased material nonlinearity and experimental variability associated with large deformations. The additional optimizer comparison in Supplementary Fig. S5 showed broadly comparable stress agreement between Adam and SSBroyden under the fixed-orientation configuration, indicating that the overall stress-prediction behavior was not strongly dependent on the optimization strategy within this exploratory analysis. Nevertheless, the model consistently preserved the overall shape and magnitude of the experimental stress-stretch responses, supporting the use of physics-informed constitutive learning for estimating myocardial mechanical behavior across different stages of RV remodeling.

### 4.2 PINN-derived biomechanical descriptors identified remodeling-associated tissue changes

Beyond stress prediction, the proposed framework simultaneously inferred specimen-specific constitutive and orientation parameters that provided an interpretable description of myocardial mechanical behavior. The constitutive parameter analysis demonstrated significant disease-associated differences in several inferred parameters (Fig. 6). In particular, the isotropic stiffness parameter *a*, fiber stiffness parameter *a*_*f*_, and cross-fiber stiffness parameter *a*_*s*_ differed significantly between disease groups, whereas the fiber-cross-fiber interaction parameter *a*_*fs*_ showed greater overlap. These findings indicate that progressive RV remodeling was associated with changes in multiple components of the inferred constitutive response rather than a uniform alteration in a single stiffness parameter.

Because model supervision was provided by the ex-vivo biaxial stress-stretch response, the inferred constitutive parameters should be interpreted as effective tissue-level descriptors of the measured passive mechanical behavior rather than as direct estimates of in-vivo myocardial material properties. Whether these inferred parameters reproduce ventricular mechanics under physiological loading remains to be established using independent in-vivo deformation measurements and organ-level mechanical validation. Given the analogous functional forms of the fiber and cross-fiber terms, these parameters are most appropriately interpreted as effective biomechanical descriptors within the reduced constitutive formulation.

The PINN-inferred orientation descriptors also exhibited remodeling-associated differences (Supplementary Fig. S4). The inferred in-plane fiber-orientation parameter *θ*_*f*_ was constrained within a global *±*20° range guided by the central distribution of histologically measured myofiber orientations across the cohort (Supplementary Fig. S4 c,d). This histological information was used only to define the physiologically plausible orientation range and was not provided to the network on a specimen- or group-specific basis. Accordingly, disease-associated differences in the inferred in-plane fiber-orientation parameter represent model-derived biomechanical trends rather than direct reconstructions of the measured histological orientations.

Collectively, the constitutive and orientation results suggest that RV remodeling is reflected through coordinated changes in tissue stiffness and in-plane material organization. These PINN-derived biomechanical descriptors therefore provided a compact mechanics-informed representation of remodeling derived from the organ-level physiological measurements used for disease classification.

### 4.3 Combining organ- and tissue-level features improved disease stratification

Integrating PINN-derived tissue-level biomechanical parameters with organ-level physiological measurements improved discrimination between control, mild PH, and severe PH compared with using organ-level measurements alone. Although the organ-level model achieved high overall classification performance, most errors occurred between the control and mild PH groups, whereas severe PH specimens were consistently identified (Fig. 7a). After augmenting the feature set with PINN-derived constitutive and orientation parameters, classification performance improved across the evaluation metrics, and only one mild PH specimen remained misclassified as control (Fig. 7b and Table 2). The largest improvement was observed for the mild PH group, indicating that the inferred tissue-level biomechanical parameters improved discrimination of the intermediate remodeling state.

Visualization of the feature representations further supported these findings. While t-SNE projections suggested improved visual separation after incorporation of the tissue-level parameters (Fig. 7c-d), pairwise centroid distances showed only modest changes between disease groups (Fig. 7e). Thus, the improvement in classification performance was more pronounced than the change in global centroid separation, indicating that the improved classification was not accompanied by a large shift in the overall group centroids.

Feature-importance analysis showed that conventional RV measurements, including RVFW thickness, RVSP, PAT, and RVEDP, remained among the influential predictors of disease stage (Fig. 8a). After incorporation of the PINN-derived parameters, the inferred in-plane fiber-orientation parameter and constitutive stiffness parameters, including *a*_*f*_ and *a*_*s*_, ranked among the most influential features together with conventional physiological measurements (Fig. 8b). These findings suggest that the PINN provided a mechanics-informed nonlinear representation of the organ-level measurements, transforming them into constitutive and orientation descriptors that improved discrimination of progressive RV remodeling when combined with the original physiological features.

### 4.4 Tissue-level mechanics enhanced characterization of intermediate RV remodeling

Mild PH represented the most challenging group to distinguish using organ-level measurements alone, with most classification errors occurring between the control and mild PH groups (Fig. 7a). This observation is consistent with mild PH representing an intermediate stage of RV remodeling, in which physiological alterations are present but less pronounced than in advanced disease. Consequently, organ-level hemodynamic and geometric measurements alone may provide limited discrimination of this intermediate remodeling state. After incorporation of the PINN-derived biomechanical descriptors, misclassification of mild PH specimens was substantially reduced, while t-SNE visualization suggested improved local separation between control and mild PH samples (Fig. 7). Pairwise centroid distances showed only modest changes, indicating that the improvement in classification performance was more pronounced than the change in global group separation. These findings suggest that the PINN-derived biomechanical descriptors provided a transformed, mechanics-informed representation of the organ-level measurements that improved discrimination of intermediate remodeling states. Because these descriptors are inferred from the physiological inputs rather than acquired as independent measurements, the observed classification improvement should be interpreted as a benefit of physics-informed feature transformation rather than the introduction of independent information. Combining these representations may therefore provide a more comprehensive characterization of disease progression, although validation in larger and longitudinal cohorts will be required to determine their utility for monitoring remodeling over time.

### 4.5 Physics-informed constitutive learning provides an alternative to conventional material parameter estimation

Conventional myocardial constitutive characterization typically relies on inverse optimization of ex-vivo mechanical experiments, in which material parameters are estimated by directly fitting constitutive model predictions to experimentally measured stress–stretch responses. This approach provides specimen-specific parameter estimates but requires mechanical testing and repeated parameter fitting for each new sample, limiting its applicability beyond experimental settings. In the present study, conventional inverse optimization also provided a reference for estimating the in-plane fiber-orientation parameter (Supplementary Fig. S4).

The proposed framework provides a different strategy by learning the relationship between organ-level physiological measurements and specimen-specific constitutive behavior during training. Once trained, the network requires only hemodynamic, functional, and geometric measurements to infer the constitutive and orientation parameters for a new specimen, without requiring specimen-specific ex-vivo parameter fitting. The corresponding stress-stretch response can then be generated by evaluating the embedded constitutive model over a prescribed deformation range. Thus, constitutive parameter inference and stress prediction are performed within a single physics-informed framework, with the inferred parameters remaining directly linked to the predicted mechanical response through the embedded constitutive formulation.

This distinction is important because the conventional inverse approach uses the measured ex-vivo mechanical response of each specimen directly during parameter estimation, whereas the PINN infers the corresponding biomechanical descriptors from organ-level measurements after learning this relationship from the training cohort. Although the present framework was developed and validated using ex-vivo biaxial mechanical data from PH, the underlying methodology is not restricted to this application. As methods for estimating myocardial deformation from clinical imaging continue to advance, similar physics-informed frameworks could potentially integrate physiological measurements with constitutive modeling to support biomechanics-informed characterization of myocardial remodeling across a broader range of SHDs.

### 4.6 Limitations and future directions

While the presented PINN framework offers significant advances in modeling myocardial mechanics, several limitations and future opportunities merit discussion:

#### (i) Dataset size and species generalizability

The proposed framework was developed using a relatively small experimental dataset obtained from rodent models of PH. The effective number of independent biological samples was therefore 39, irrespective of the multiple stress-stretch measurements available for each specimen. Although LOOCV maximized data utilization, the limited cohort size, particularly for the mild PH group, reduced statistical power and may not fully represent the biological variability associated with RV remodeling. Future studies including larger experimental datasets and independent human cohorts will be important for establishing the robustness, reproducibility, and clinical generalizability of the proposed framework.

#### (ii) Ex-vivo mechanical supervision

The present framework was trained using ex-vivo biaxial mechanical testing, where experimentally measured stresses were available to supervise PINN training. Accordingly, agreement with the measured ex-vivo stress-stretch response does not by itself establish that the inferred constitutive parameters reproduce myocardial mechanics under in-vivo physiological loading. Future validation should therefore couple the inferred constitutive parameters with subject-specific ventricular models and independent in-vivo deformation or loading measurements to evaluate their ability to reproduce organ-level cardiac mechanics. Although ex-vivo testing provides a controlled setting for evaluating the proposed methodology, future clinical translation will require further validation using medical imaging or other clinically accessible measurements while preserving the underlying constitutive constraints.

#### (iii) Reduced constitutive parameterization

To improve parameter identifiability, the exponential constitutive coefficients (*b, b*_*f*_, *b*_*s*_, *b*_*fs*_) were fixed at *b* = *b*_*f*_ = *b*_*s*_ = *b*_*fs*_ = 1 across all specimens during training, while the constitutive magnitude parameters and in-plane fiber-orientation parameter were inferred. The reduced formulation imposes orthogonality between the in-plane material directions and restricts the orientation range, while the fiber and cross-fiber stiffness parameters retain analogous constitutive forms. Accordingly, interpretation of individual parameter values should be considered within the context of this reduced parameterization. Although this strategy produced a stable inverse problem for the available dataset, future studies with larger and more diverse datasets may enable reliable estimation of a broader constitutive parameter set and evaluation of more complex constitutive formulations.

#### (iv) Constitutive assumptions

The present framework adopted an incompressible hyperelastic constitutive formulation, which neglects volumetric compressibility and time-dependent mechanical behavior. While these assumptions are appropriate for the quasi-static biaxial experiments considered in this study, future work may incorporate compressible and visco-hyperelastic constitutive models to better represent myocardial mechanics under more complex physiological loading conditions.

#### (v) Loading-protocol generalizability

The present framework was trained and evaluated using a single equibiaxial (1:1) loading protocol. Although the embedded constitutive formulation can be evaluated at other combinations of circumferential and longitudinal stretch, prediction under non-equibiaxial loading conditions was not experimentally validated in the present cohort. Future studies incorporating multiple biaxial loading ratios will be needed to assess the ability of the framework to generalize across a broader range of deformation states. In addition, biaxial stretch was quantified from the overall specimen deformation and therefore represents a nominal measure of tissue deformation. Future studies incorporating optical tracking within the central gauge region could provide local strain measurements and further characterize spatial deformation heterogeneity.

#### (vi) Strain and sex imbalance across disease groups

The experimental design was not fully balanced across strain, sex, and disease stage, with the mild-PH group consisting only of Sprague–Dawley animals and relatively few female animals in some subgroups. These imbalances limited the ability to assess strain- and sex-dependent differences in RV biomechanical remodeling.

#### (viii) Interpretation of inferred orientation

The inferred orientation parameter represents an effective in-plane material direction within the circumferential-longitudinal plane. The orthogonal cross-fiber direction used in the reduced constitutive formulation is distinct from the anatomical myocardial sheet direction associated with the three-dimensional laminar architecture. Histological measurements were used only to define the global physiologically plausible range for the fiber-orientation parameter and were not incorporated as specimen-specific supervision. Accordingly, the inferred orientation should be interpreted as a biomechanical descriptor of in-plane tissue organization rather than as a direct reconstruction of the full three-dimensional myocardial microstructure. Future studies combining spatially resolved histology, additional loading protocols, and three-dimensional structural measurements may enable more detailed validation of inferred myocardial architecture.

#### (ix) In-vivo pre-strain and reference configuration

The organ-level physiological measurements were acquired under in-vivo loading, whereas the ex-vivo biaxial response was referenced to the unloaded specimen configuration used during mechanical testing. Physiological pre-strain and residual deformation were therefore not explicitly modeled and may contribute to variability in the learned relationship between organ-level measurements and tissue-level constitutive behavior. Future studies could estimate subject-specific pre-strain or unloaded reference configurations from imaging-derived deformation and incorporate this information directly into the physics-informed framework.

## Acknowledgements

This work was supported by the American Heart Association (25PRE1377496) to R.R.M., the National Institutes of Health (R56HL172052), and the National Science Foundation (2244995) to R.A.

## Declaration of competing interest

All authors declare no conflict of interest.

## Data availability

The data that support the findings of this study are available from the corresponding author, R.A., upon request.

## Notes

### Competing Interest Statement

The authors have declared no competing interest.

## References

1. Wheat, H. L., Fedson, S., Bozkurt, B. & Josephson, R. A. Cardiac rehabilitation in heart failure: Indications for exercise training based on heart failure phenotype. Prog. Cardiovasc. Dis. 70, 16–21 (2022).

2. Arts, T., Delhaas, T., Bovendeerd, P., Verbeek, X. & Prinzen, F. W. Adaptation to mechanical load determines shape and properties of heart and circulation: the circadapt model. Am. J. Physiol. Circ. Physiol. 288, H1943–H1954 (2005).

3. Mendiola, E. A., da Silva Gonçalves Bos, D., Leichter, D. M., Vang, A., Zhang, P., Leary, O. P., Gilbert, R. J., Avazmohammadi, R. & Choudhary, G. Right ventricular architectural remodeling and functional adaptation in pulmonary hypertension. Circ. Hear. Fail. 16, e009768 (2023).

4. Neelakantan, S., Kumar, M., Mendiola, E. A., Phelan, H., Serpooshan, V., Sadayappan, S. & Avazmohammadi, R. Multiscale characterization of left ventricle active behavior in the mouse. Acta biomaterialia 162, 240–253 (2023).

5. Neelakantan, S., Vang, A., Mehdi, R. R., Phelan, H., Nicely, P., Imran, T., Zhang, P., Choudhary, G. & Avazmohammadi, R. Right ventricular stiffening and anisotropy alterations in pulmonary hypertension: mechanisms and relations to right heart failure. J. Am. Hear. Assoc. 14, e037126 (2025).

6. Usman, M., Mendiola, E. A., Mukherjee, T., Mehdi, R. R., Ohayon, J., Alluri, P. G., Sadayappan, S., Choudhary, G. & Avazmohammadi, R. On the possibility of estimating myocardial fiber architecture from cardiac strains. In International Conference on Functional Imaging and Modeling of the Heart, 74–83 (Springer, 2023).

7. Nguyen-Truong, M. & Wang, Z. Biomechanical properties and mechanobiology of cardiac ecm. Cardiac Extracell. Matrix: Fundamental Sci. to Clin. Appl. 1–19 (2018).

8. Lai, Y.-C., Potoka, K. C., Champion, H. C., Mora, A. L. & Gladwin, M. T. Pulmonary arterial hypertension: the clinical syndrome. Circ. research 115, 115–130 (2014).

9. Simonneau, G., Gatzoulis, M. A., Adatia, I., Celermajer, D., Denton, C., Ghofrani, A., Gomez Sanchez, M. A., Krishna Kumar, R., Landzberg, M., Machado, R. F. et al. Updated clinical classification of pulmonary hypertension. J. Am. Coll. Cardiol. 62, D34–D41 (2013).

10. Humbert, M., Guignabert, C., Bonnet, S., Dorfmüller, P., Klinger, J. R., Nicolls, M. R., Olschewski, A. J., Pullamsetti, S. S., Schermuly, R. T., Stenmark, K. R. et al. Pathology and pathobiology of pulmonary hypertension: state of the art and research perspectives. Eur. Respir. J. 53 (2019).

11. Neelakantan, S., Mendiola, E. A., Zambrano, B., Vang, A., Myers, K. J., Zhang, P., Choudhary, G. & Avazmohammadi, R. Dissecting contributions of pulmonary arterial remodeling to right ventricular afterload in pulmonary hypertension. Bioeng. & Transl. Medicine 10, e70035 (2025).

12. Noordegraaf, A. V., Chin, K. M., Haddad, F., Hassoun, P. M., Hemnes, A. R., Hopkins, S. R., Kawut, S. M., Langleben, D., Lumens, J. & Naeije, R. Pathophysiology of the right ventricle and of the pulmonary circulation in pulmonary hypertension: an update. Eur. Respir. J. 53 (2019).

13. Van Der Bruggen, C. E., Tedford, R. J., Handoko, M. L., Van Der Velden, J. & de Man, F. S. Rv pressure overload: from hypertrophy to failure. Cardiovasc. research 113, 1423–1432 (2017).

14. Voelkel, N. F., Quaife, R. A., Leinwand, L. A., Barst, R. J., McGoon, M. D., Meldrum, D. R., Dupuis, J., Long, C. S., Rubin, L. J., Smart, F. W. et al. Right ventricular function and failure: report of a national heart, lung, and blood institute working group on cellular and molecular mechanisms of right heart failure. Circulation 114, 1883–1891 (2006).

15. Bernardo, R. J., Haddad, F., Couture, E. J., Hansmann, G., de Jesus Perez, V.A., Denault, A. Y., de Man, F. S. & Amsallem, M. Mechanics of right ventricular dysfunction in pulmonary arterial hypertension and heart failure with preserved ejection fraction. Cardiovasc. Diagn. Ther. 10, 1580 (2020).

16. Raymond, R. J., Hinderliter, A. L., Willis, P. W., Ralph, D., Caldwell, E. J., Williams, W., Ettinger, N. A., Hill, N. S., Summer, W. R., de Boisblanc, B. et al. Echocardiographic predictors of adverse outcomes in primary pulmonary hypertension. J. Am. Coll. Cardiol. 39, 1214–1219 (2002).

17. Haddad, F., Doyle, R., Murphy, D. J. & Hunt, S. A. Right ventricular function in cardiovascular disease, part ii: pathophysiology, clinical importance, and management of right ventricular failure. Circulation 117, 1717–1731 (2008).

18. Ghio, S., Klersy, C., Magrini, G., D’Armini, A. M., Scelsi, L., Raineri, C., Pasotti, M., Serio, A., Campana, C. & Viganò, M. Prognostic relevance of the echocardiographic assessment of right ventricular function in patients with idiopathic pulmonary arterial hypertension. Int. journal cardiology 140, 272–278 (2010).

19. Purmah, Y., Lei, L. Y., Dykstra, S., Mikami, Y., Cornhill, A., Satriano, A., Flewitt, J., Rivest, S., Sandonato, R., Seib, M. et al. Right ventricular ejection fraction for the prediction of major adverse cardiovascular and heart failure-related events: a cardiac mri based study of 7131 patients with known or suspected cardiovascular disease. Circ. Cardiovasc. Imaging 14, e011337 (2021).

20. Fine, N. M., Chen, L., Bastiansen, P. M., Frantz, R. P., Pellikka, P. A., Oh, J. K. & Kane, G. C. Outcome prediction by quantitative right ventricular function assessment in 575 subjects evaluated for pulmonary hypertension. Circ. Cardiovasc. Imaging 6, 711–721 (2013).

21. Holzapfel, G. A. & Ogden, R. W. Constitutive modelling of passive myocardium: a structurally based framework for material characterization. Philos. Transactions Royal Soc. A: Math. Phys. Eng. Sci. 367, 3445–3475 (2009).

22. Mehdi, R. R., Neelakantan, S., Wang, E., Zhang, P., Choudhary, G. & Avazmohammadi, R. Abstract p2008: Contractile adaptation of the right ventricular myocardium in pulmonary hypertension. Circ. Res. 133, AP2008–AP2008 (2023).

23. Emig, R., Zgierski-Johnston, C. M., Timmermann, V., Taberner, A. J., Nash, M. P., Kohl, P. & Peyronnet, R. Passive myocardial mechanical properties: meaning, measurement, models. Biophys. reviews 1–24 (2021).

24. Li, D. S., Mendiola, E. A., Avazmohammadi, R., Sachse, F. B. & Sacks, M. S. A multi-scale computational model for the passive mechanical behavior of right ventricular myocardium. J. mechanical behavior biomedical materials 142, 105788 (2023).

25. Taç, V., Rausch, M. K., Costabal, F. S. & Tepole, A. B. Data-driven anisotropic finite viscoelasticity using neural ordinary differential equations. Comput. methods applied mechanics engineering 411, 116046 (2023).

26. Mehdi, R. R., Kadivar, N., Mukherjee, T., Mendiola, E. A., Bersali, A., Shah, D. J., Karniadakis, G. & Avazmohammadi, R. Non-invasive diagnosis of chronic myocardial infarction via composite in-silico-human data learning. Adv. Sci. e06933 (2025).

27. Mehdi, R. R., Kadivar, N., Serpooshan, V., Myers, K. J., Karniadakis, G. & Avazmohammadi, R. Towards non-invasive estimation of myocardial scar stiffness from cardiac strains using deep learning. In International Conference on Functional Imaging and Modeling of the Heart, 420–429 (Springer, 2025).

28. Mehdi, R. R., Mendiola, E. A., Sears, A., Choudhary, G., Ohayon, J., Pettigrew, R. & Avazmohammadi, R. Comparison of three machine learning methods to estimate myocardial stiffness. In Reduced Order Models for the Biomechanics of Living Organs, 363–382 (Elsevier, 2023).

29. Karniadakis, G. E., Kevrekidis, I. G., Lu, L., Perdikaris, P., Wang, S. & Yang, L. Physics-informed machine learning. Nat. Rev. Phys. 3, 422–440 (2021).

30. Song, S. & Jin, H. Identifying constitutive parameters for complex hyperelastic materials using physics-informed neural networks. Soft Matter 20, 5915–5926 (2024).

31. Caforio, F., Regazzoni, F., Pagani, S., Karabelas, E., Augustin, C., Haase, G., Plank, G. & Quarteroni, A. Physics-informed neural network estimation of material properties in soft tissue nonlinear biomechanical models. Comput. Mech. 1–27 (2024).

32. Holzapfel, G. A., Linka, K., Sherifova, S. & Cyron, C. J. Predictive constitutive modelling of arteries by deep learning. J. Royal Soc. Interface 18, 20210411 (2021).

33. Linka, K., Cavinato, C., Humphrey, J. D. & Cyron, C. J. Predicting and understanding arterial elasticity from key microstructural features by bidirectional deep learning. Acta biomaterialia 147, 63–72 (2022).

34. Buoso, S., Joyce, T. & Kozerke, S. Personalising left-ventricular biophysical models of the heart using parametric physics-informed neural networks. Med. Image Analysis 71, 102066 (2021).

35. Sahli Costabal, F., Yang, Y., Perdikaris, P., Hurtado, D. E. & Kuhl, E. Physics-informed neural networks for cardiac activation mapping. Front. Phys. 8, 42 (2020).

