## Supplementary file for "Physics-informed neural network for predicting myocardial mechanics from in vivo physiological measurements"

### SUPPLEMENTAL MATERIAL

### Data S1. Supplemental Methods

Additional experimental details for the in-vivo measurements, biaxial mechanical testing, and histological preparation were adapted from previously reported protocols[1] and are summarized below for completeness.

#### Data S1.1 In-vivo physiological measurements and additional cardiac characterization

At the terminal study time point, animals were maintained on a heating pad at approximately 37 °C and anesthetized using continuous inhaled isoflurane during echocardiographic and catheterization measurements. Transthoracic echocardiography was performed using a high-frequency linear-array transducer and a Vevo 2100 imaging system. Pulmonary acceleration time was obtained from pulsed-wave Doppler recordings at the right ventricular (RV) outflow tract. Tricuspid annular plane systolic excursion was measured from M-mode recordings as the systolic displacement of the tricuspid annulus. Tissue Doppler imaging was used to quantify RV systolic myocardial velocity and additional diastolic myocardial and valvular inflow velocities. Echocardiographic measurements were analyzed in a blinded manner.

Following echocardiography, invasive RV catheterization was performed using a high-fidelity pressure catheter introduced through the RV apex. Right ventricular systolic pressure and right ventricular end-diastolic pressure were recorded to characterize RV pressure loading and diastolic function. These measurements, together with the echocardiographic functional and geometric indices, were used to characterize disease severity and constituted the organ-level physiological input feature set for the proposed physics-informed neural network (PINN).

Additional echocardiographic measurements were acquired for broader characterization of cardiac function, including diastolic myocardial velocities, tricuspid and mitral inflow velocities, and left ventricular outflow tract dimensions and Doppler-derived hemodynamic indices. These measurements are summarized in Supplementary Table S1 and were not included in the final PINN input feature set.

#### Data S1.2 Biaxial mechanical testing

Following terminal physiological measurements, the RV free-wall (RVFW) was isolated and trimmed into rectangular specimens aligned with the circumferential and longitudinal directions. Specimen thickness was measured using calipers before mechanical testing. Each specimen was mounted in a CellScale BioTester (CellScale, Waterloo, Ontario, Canada) using BioRakes, with four metal pins on each rake providing multi-point tissue attachment. The specimens were submerged in phosphate-buffered saline at approximately 22°C during testing to prevent tissue dehydration.

No optical markers were used for non-contact displacement tracking. The specimens underwent 10 preconditioning cycles followed by 10 equibiaxial loading cycles, with both the circumferential and longitudinal directions stretched by approximately 30%. The unloaded specimen configuration at the beginning of the recorded loading cycle was taken as the reference configuration for calculation of stretch.

A specimen-level nominal stretch was defined as the ratio of the current specimen length to its reference length,

$$\lambda = \frac{L}{L_0}, \tag{1}$$

where $L$ and $L_0$ denote the current and reference specimen lengths, respectively. This definition provides a specimen-level measure of deformation based on the overall change in specimen length; local strain fields within the central gauge region were not quantified.

The measured specimen dimensions were used to determine the initial cross-sectional areas in the circumferential and longitudinal directions. Force measurements were converted into first Piola-Kirchhoff stresses according to

$$P_{\text{circ}} = \frac{F_{\text{circ}}}{A_{0,\text{circ}}}, \qquad P_{\text{long}} = \frac{F_{\text{long}}}{A_{0,\text{long}}}, \tag{2}$$

where $F_{\text{circ}}$ and $F_{\text{long}}$ are the measured forces and $A_{0,\text{circ}}$ and $A_{0,\text{long}}$ are the corresponding initial cross-sectional areas. The resulting circumferential and longitudinal stress-stretch responses were used as the experimental targets for PINN training and evaluation.

For selected analyses, total stress was defined as the sum of the two directional stress components,

$$P_{\text{tot}} = P_{\text{circ}} + P_{\text{long}}. \tag{3}$$

#### Data S1.3 Histological preparation and imaging

After biaxial mechanical testing, RVFW specimens were fixed in 10% neutral buffered formalin and subsequently stored in 70% ethanol. The specimens were sectioned through the wall thickness at a thickness of approximately 5 $\mu$m and stained using picrosirius red to characterize myocardial and collagen organization.

The stained sections were mounted on glass slides and imaged using a whole-slide scanning microscope at high magnification. Histological analysis was used to characterize the distribution of myofiber and collagen orientations across the ventricular wall, as summarized in Supplementary Fig. S4. The central distribution of histologically measured myofiber orientations across the cohort was used to define the global $\pm 20^\circ$ range for the PINN-inferred orientation parameter $\theta_f$. Because $\theta_f$ represents an effective in-plane material orientation within the reduced biaxial constitutive formulation rather than a direct reconstruction of local transmural microstructure, specimen-specific histological orientations were not imposed as constitutive directions or used as PINN inputs or supervision targets during model training.

#### Data S1.4 Comparison of first- and second-order optimization strategies

The results presented in the main manuscript were obtained using the final PINN framework trained with the Adam optimizer, in which the in-plane fiber-orientation parameter $\theta_f$ was inferred separately for each specimen within the prescribed $\pm 20^\circ$ range. Figure S2 provides additional evaluation of this model by summarizing the specimen-wise distributions of the coefficient of determination ($R^2$) together with Bland-Altman analyses comparing the predicted and experimentally measured total stresses at representative mid- and maximum-stretch levels. Prediction differences were smaller at the representative intermediate stretch than at the maximum stretch, indicating closer agreement at the intermediate loading level.

To examine the influence of optimization strategy independently of orientation estimation, an exploratory comparison was performed between the first-order Adam optimizer and the self-scaled second-order optimizer (SSBroyden)[2]. For this analysis, the in-plane fiber-orientation parameter was fixed at $\theta_f = -3^\circ$ for all specimens, with the orthogonal cross-fiber direction defined accordingly. The fixed orientation was selected because the cohort-averaged histological myofiber orientation was close to $-3^\circ$ (Supplementary Fig. S4). This fixed-orientation analysis was performed separately from the main PINN framework, in which $\theta_f$ was inferred for each specimen within the global $\pm 20^\circ$ range defined from the cohort-level histological measurements; specimen-specific histological orientations were not imposed. The same fixed-orientation configuration was used for both optimization strategies to ensure a consistent comparison.

Training was formulated as the optimization problem

$$\min_{\theta} \mathscr{L}(\theta), \tag{4}$$

where $\theta$ denotes the trainable network parameters and $\mathscr{L}$ is the total physics-informed loss comprising the stress, reference, monotonicity, and regularization terms described in the main manuscript.

For the Adam optimizer, the network parameters were updated according to

$$\theta_{k+1} = \theta_k - \alpha_k \nabla \mathscr{L}(\theta_k), \tag{5}$$

where $\alpha_k$ denotes the adaptive learning rate. In contrast, SSBroyden incorporates curvature information through an evolving approximation of the inverse Hessian,

$$\theta_{k+1} = \theta_k - \mathbf{H}_k^{-1} \nabla \mathscr{L}(\theta_k), \tag{6}$$

where $\mathbf{H}_k^{-1}$ denotes the approximate inverse Hessian matrix.

Supplementary Fig. S5 compares the agreement between predicted and experimentally measured total stresses obtained using the two optimization strategies under the fixed-orientation configuration. Overall prediction performance was comparable between Adam and SSBroyden. At the intermediate stretch level, SSBroyden exhibited a slightly narrower range of prediction differences than Adam, whereas similar agreement was observed near the maximum stretch. These results indicate that the overall stress-prediction behavior was not strongly dependent on the optimization strategy considered under the fixed-orientation configuration.

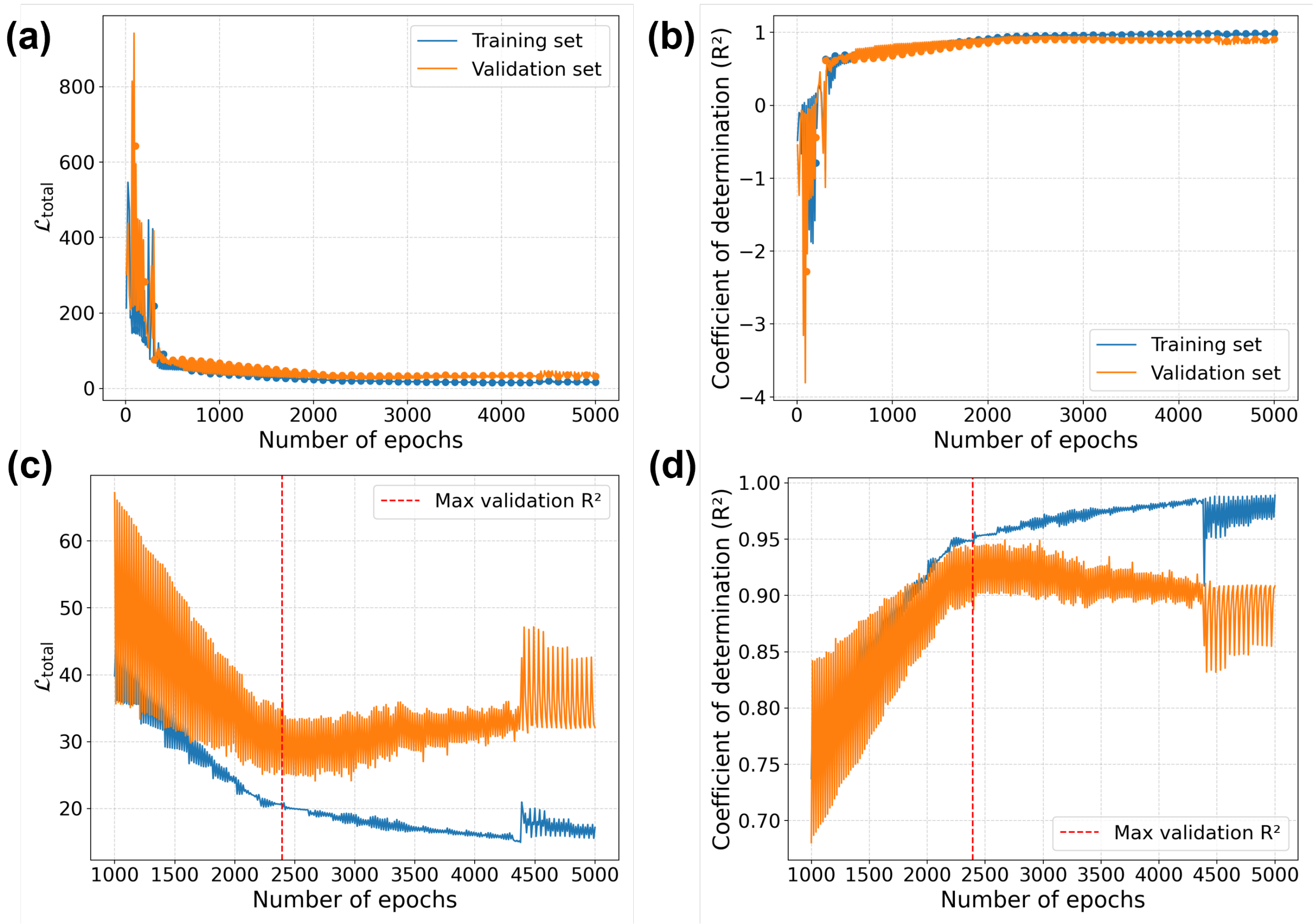


**Fig. S1.** Training dynamics of the physics-informed neural network (PINN). (a) Training and validation total loss as a function of training epochs. (b) Corresponding coefficient of determination ($R^2$) for the training and validation sets. (c,d) Enlarged views of the late-stage training period highlighting convergence of the total loss and $R^2$, respectively. The model corresponding to the minimum validation loss was retained for subsequent evaluation.

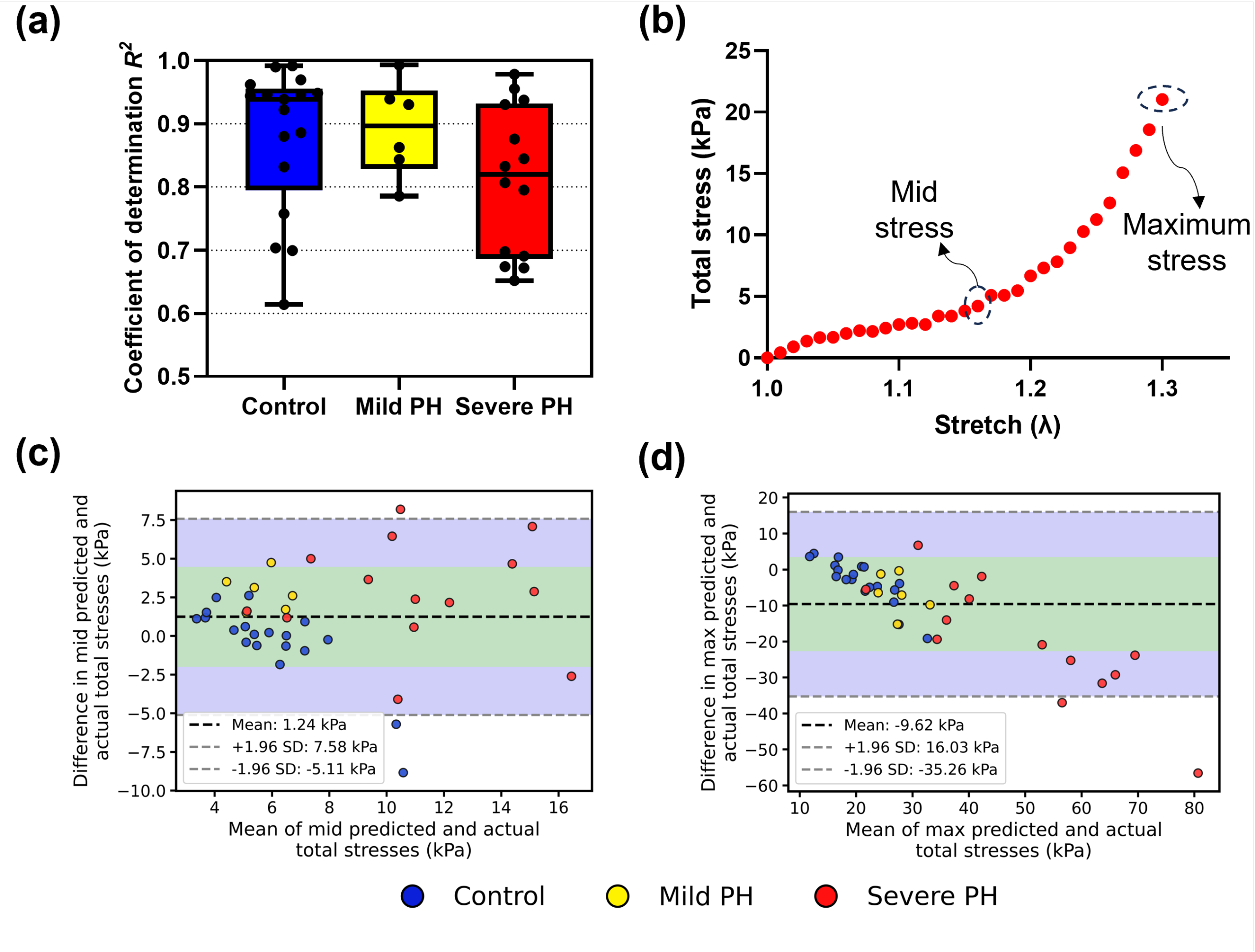


**Fig. S2.** Additional evaluation of PINN stress-prediction performance. (a) Distribution of the coefficient of determination ($R^2$) across the control, mild pulmonary hypertension (PH), and severe PH groups. (b) Representative total stress-stretch response illustrating the mid- and maximum-stretch locations used for quantitative comparison. (c,d) Bland-Altman analyses comparing predicted and experimentally measured total stresses at the representative mid- and maximum-stretch levels, respectively, for models trained using the Adam optimizer.

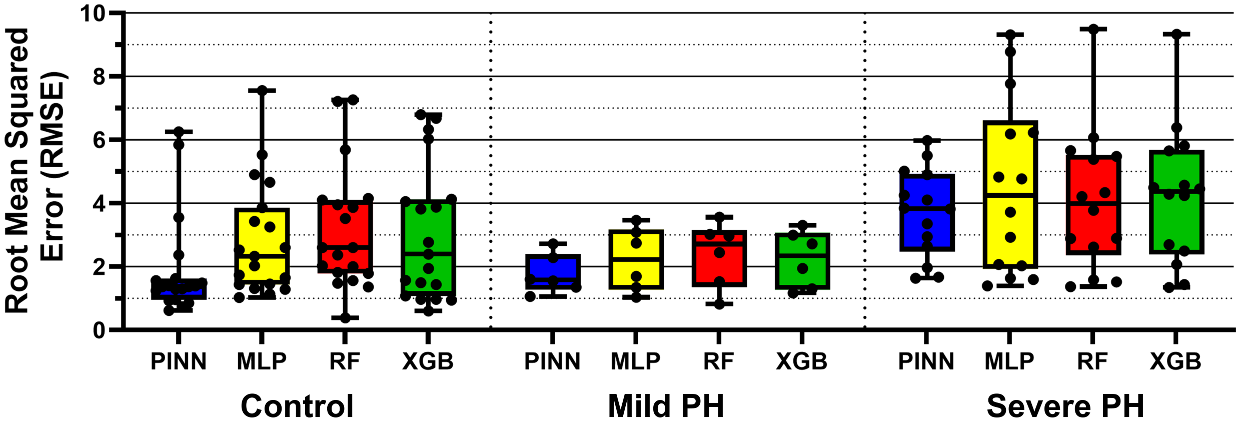


**Fig. S3.** Comparison of root mean squared error (RMSE) between the proposed physics-informed neural network (PINN) and conventional machine-learning regression models. Specimen-wise RMSE values are shown separately for the control, mild pulmonary hypertension (PH), and severe PH groups for the PINN, multilayer perceptron (MLP), random forest (RF), and extreme gradient boosting (XGBoost) models. Each point represents an individual specimen, boxes indicate the interquartile range with the median shown by the horizontal line, and whiskers indicate the range of the observed values.

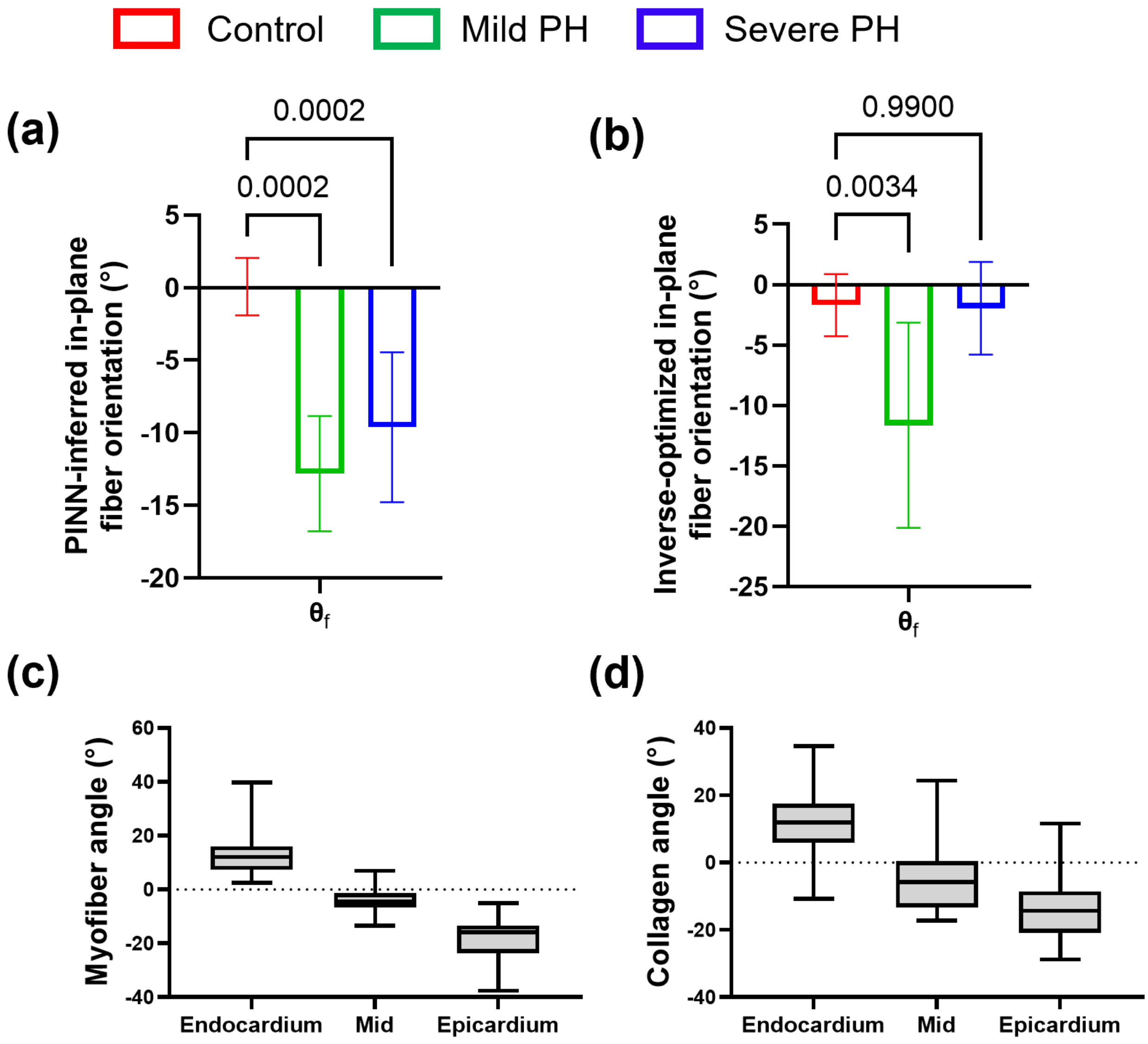


**Fig. S4.** Comparison of PINN-inferred orientation descriptors and histologically measured myocardial orientations. (a) PINN-inferred in-plane fiber-orientation parameter ($\theta_f$) across the control, mild pulmonary hypertension (PH), and severe PH groups. (b) Corresponding in-plane fiber-orientation estimates obtained using conventional inverse optimization. (c) Histologically measured myofiber orientations across the endocardial, mid-wall, and epicardial regions. (d) Histologically measured collagen orientations across the corresponding transmural regions. Error bars in (a,b) represent the 95% confidence intervals across specimens within each disease group. Statistical comparisons in (a,b) were performed using ordinary one-way analysis of variance (ANOVA) followed by Dunnett's multiple-comparisons test, with the control group as the reference; adjusted $p$-values are indicated above the corresponding panels.

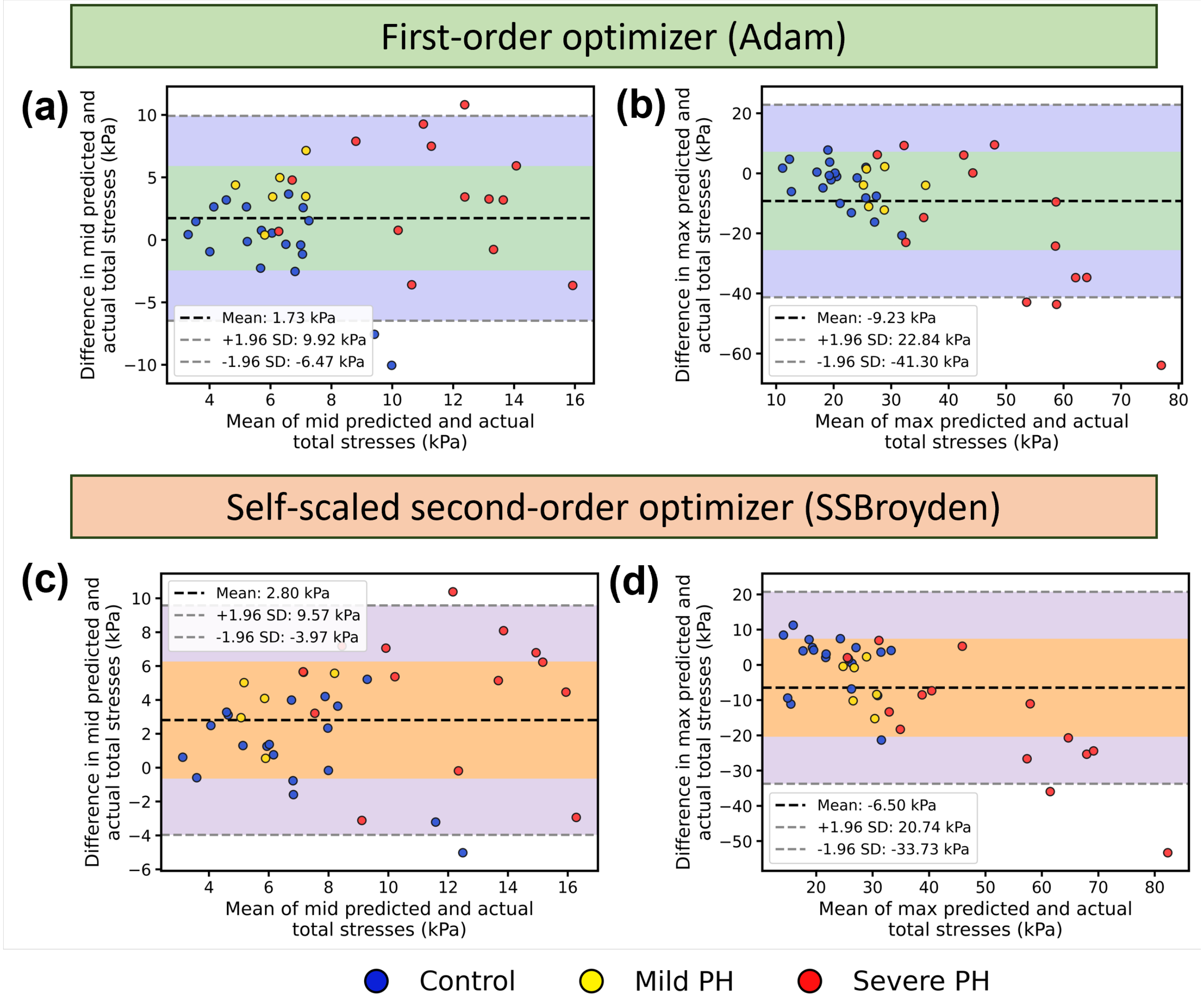


**Fig. S5.** Comparison of first-order and second-order optimization strategies for PINN training. Bland-Altman analyses comparing predicted and experimentally measured total stresses at the (a,c) mid-stretch and (b,d) maximum-stretch levels for models trained using the first-order Adam optimizer (top row) and the self-scaled second-order optimizer (SSBroyden) (bottom row). Dashed lines indicate the mean difference and the 95% limits of agreement.

**Table S1.** Additional echocardiographic parameters for control and pulmonary hypertension (PH) groups. Values are reported as mean ± standard deviation. Listed measures include tissue Doppler indices (late and early diastolic myocardial velocities A′/E′, tricuspid and mitral inflow velocities (A and E waves), and tricuspid valve lateral wall velocities (TV LW A′/E′), which together characterize diastolic and systolic function. Left ventricular outflow tract (LVOT) dimensions and Doppler-derived indices (length, velocity-time integral, mean and peak velocities, and corresponding pressure gradients) are also reported. These parameters provide a complementary assessment of global cardiac performance and reveal differences between control and PH groups beyond standard metrics. Control and PH groups were compared using a two-tailed unpaired Welch's $t$-test.

| Diastolic function | | | |
|---|---|---|---|
| | Control | PH | $p$-value |
| Late diastolic myocardial velocity ($A'$, mm/s) | -44.47 ± 13.60 | -39.34 ± 15.84 | 0.3514 |
| Early diastolic myocardial velocity ($E'$, mm/s) | -39.95 ± 10.89 | -23.01 ± 10.67 | 0.0001 |
| Tricuspid valve late diastolic inflow velocity (TV A, mm/s) | 602.11 ± 150.41 | 521.07 ± 245.45 | 0.2598 |
| Tricuspid valve early diastolic inflow velocity (TV E, mm/s) | 346.33 ± 110.92 | 484.56 ± 145.77 | 0.0208 |
| Tricuspid valve lateral wall late diastolic velocity (TV LW $A'$, mm/s) | -35.93 ± 17.11 | -31.33 ± 14.23 | 0.2988 |
| Tricuspid valve lateral wall early diastolic velocity (TV LW $E'$, mm/s) | -29.38 ± 11.58 | -22.63 ± 6.18 | 0.0522 |
| Mitral valve late diastolic inflow velocity (MV A, mm/s) | 616.01 ± 176.42 | 550.13 ± 185.57 | 0.2385 |
| Mitral valve early diastolic inflow velocity (MV E, mm/s) | 930.73 ± 110.60 | 615.49 ± 266.65 | 0.0008 |
| Left ventricular outflow tract (LVOT) hemodynamics | | | |
| LVOT length (mm) | 2.44 ± 0.38 | 2.40 ± 0.36 | 0.9109 |
| LVOT velocity-time integral (LVOT VTI, mm) | 50.30 ± 13.34 | 38.59 ± 15.39 | 0.0047 |
| LVOT mean velocity (mm/s) | -464.77 ± 113.88 | -372.31 ± 116.48 | 0.0044 |
| LVOT mean pressure gradient (mmHg) | 0.91 ± 0.43 | 0.61 ± 0.43 | 0.0046 |
| LVOT peak velocity (mm/s) | -997.30 ± 197.81 | -855.34 ± 223.22 | 0.0322 |
| LVOT peak pressure gradient (mmHg) | 4.16 ± 1.59 | 3.12 ± 1.69 | 0.0392 |
| LVOT stroke volume ($\mu$L) | 360.15 ± 99.15 | 169.91 ± 66.61 | 0.0083 |
| LVOT cardiac output (mL/min) | 108.23 ± 36.97 | 66.02 ± 35.96 | 0.0065 |

#### References


1. Neelakantan, S., Vang, A., Mehdi, R. R., Phelan, H., Nicely, P., Imran, T., Zhang, P., Choudhary, G. & Avazmohammadi, R. Right ventricular stiffening and anisotropy alterations in pulmonary hypertension: mechanisms and relations to right heart failure. *J. Am. Hear. Assoc.* **14**, e037126 (2025).
2. Kiyani, E., Shukla, K., Urbán, J. F., Darbon, J. & Karniadakis, G. E. Optimizing the optimizer for physics-informed neural networks and kolmogorov-arnold networks. *Comput. Methods Appl. Mech. Eng.* **446**, 118308 (2025).
